## Supplementary Figures and Tables for "Systematic benchmarking of tools for CpG methylation detection from Nanopore sequencing"

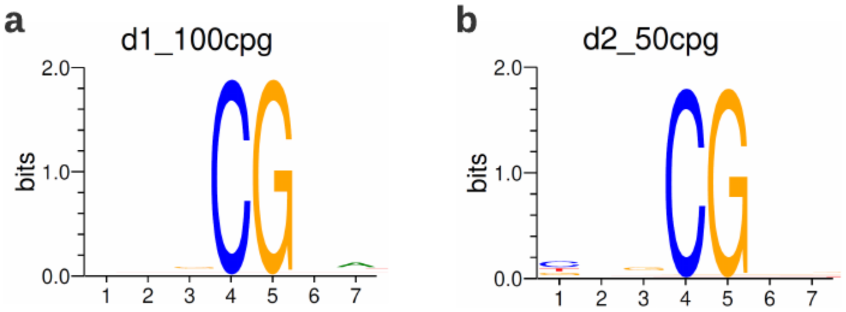


**Supplementary Figure 1. Sequence contexts of the CpG sites in datasets 1 and 2.** We show the sequence logos with the information content in bits (y axis) for the positions surrounding the CpG sites used in mixture dataset 1 **(a)** and 2 **(b).** The two datasets do not show strong differences in the sequence context around the selected sites.

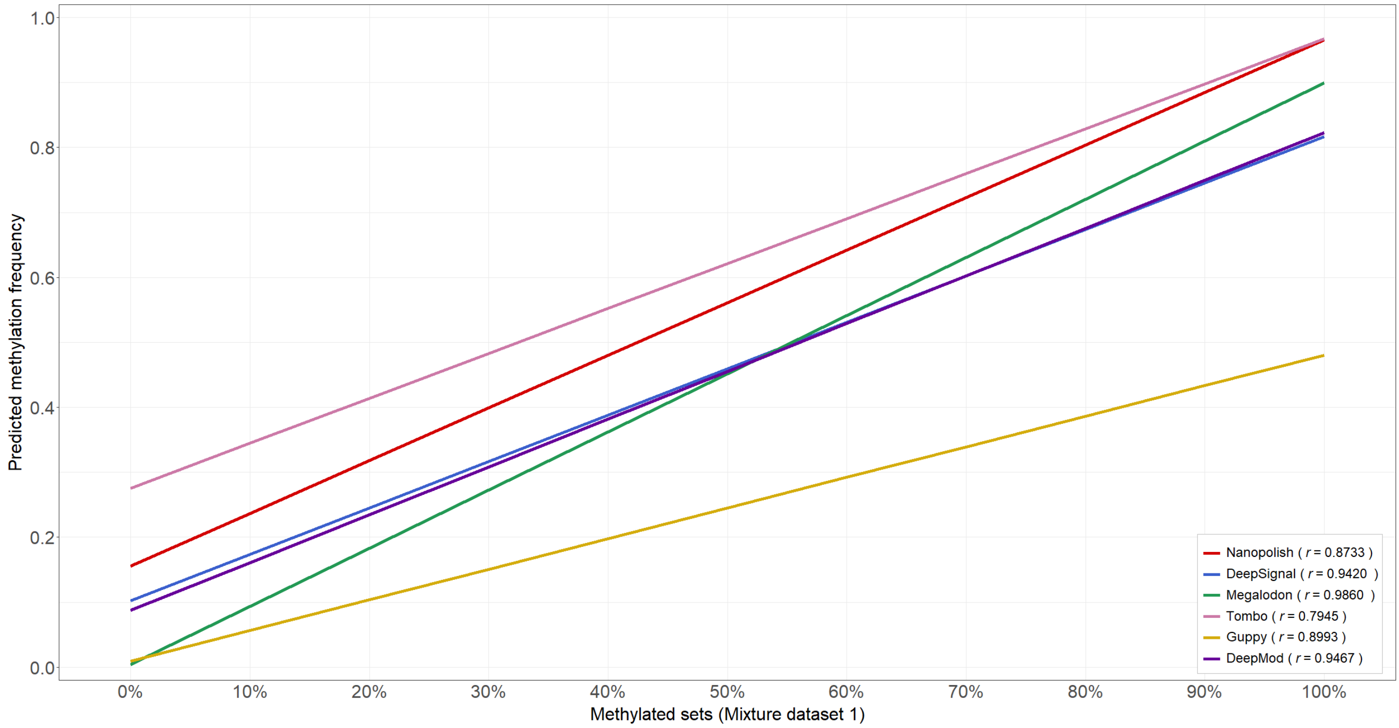


**Supplementary Figure 2. Regression lines of the predictions on the mixture dataset 1.** This plot uses the same data as Fig. 2a but only showing regression lines and the Pearson's correlation (r) for each tool.

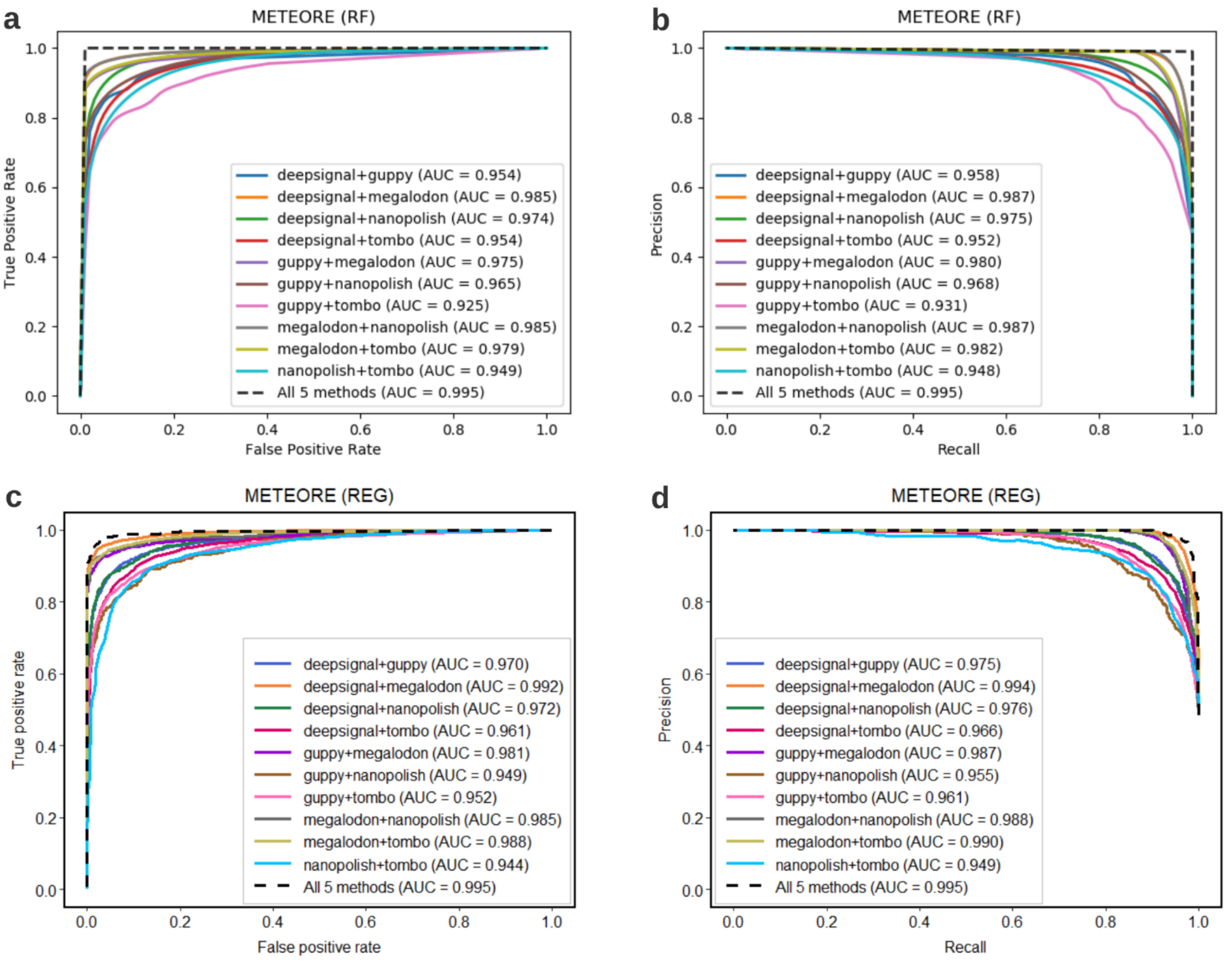


**Supplementary Figure 3. Model accuracy for METEORE at the individual read level. a** Receiver operating characteristic (ROC) curves showing the false positive rate (x axis) and true positive rate (y axis) for the predictions of the METEORE random forest (RF) model combining two methods (with default parameters: n_estimator = 10 and max_depth = None) using the sites of the mixture dataset 1 at individual read level. The curves were built from the average of a 10-fold cross validation. **b** Precision-recall (PR) curves showing the recall (x axis) and precision (y axis) for the RF model using the sites of the mixture dataset 1 at individual read level, also built from 10-fold cross validation. **c** ROC curves for the METEORE regression (REG) model combining two methods at individual read level. Curves were built from a 5-fold cross validation using the sites of mixture dataset 1. **d** PR curves for the METEORE REG model using the sites of the mixture dataset 1 at individual read level, built from a 5-fold cross validation.

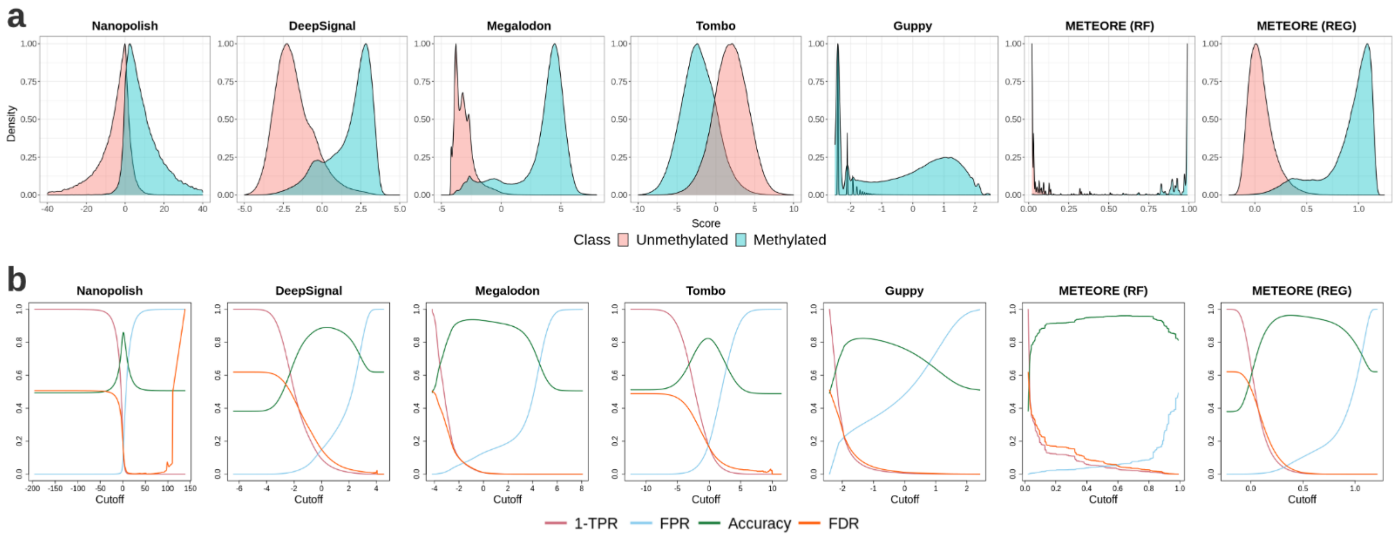


**Supplementary Figure 4. Score distributions and accuracy metrics. a** Score (x axis) distribution for methylated and unmethylated sites in individual reads for each tested tool on the mixture dataset 1, including METEORE. We used METEORE combining the predictions of Megalodon and DeepSignal with a random forest (RF) (parameters: max_dep=3 and n_estimator=10) and with a regression model (REG). **b** Distribution of various accuracy metrics (y axis) according to the score (x axis) for each method shown in **(a)**. We show the false positive rate (FPR), false discovery rate (FDR), 1 - true positive rate (TPR), and Accuracy curves as a function of the single score (x axis) cutoff for each tool, where FPR = FP/(FP+TN), TPR = TP/(TP+FN), FDR = FP/(TP+FP), accuracy = (TP + TN)/(TP + TN + FP + FN); and TP = true positives, FP = false positives, TN = true negatives, and FN = false negatives.

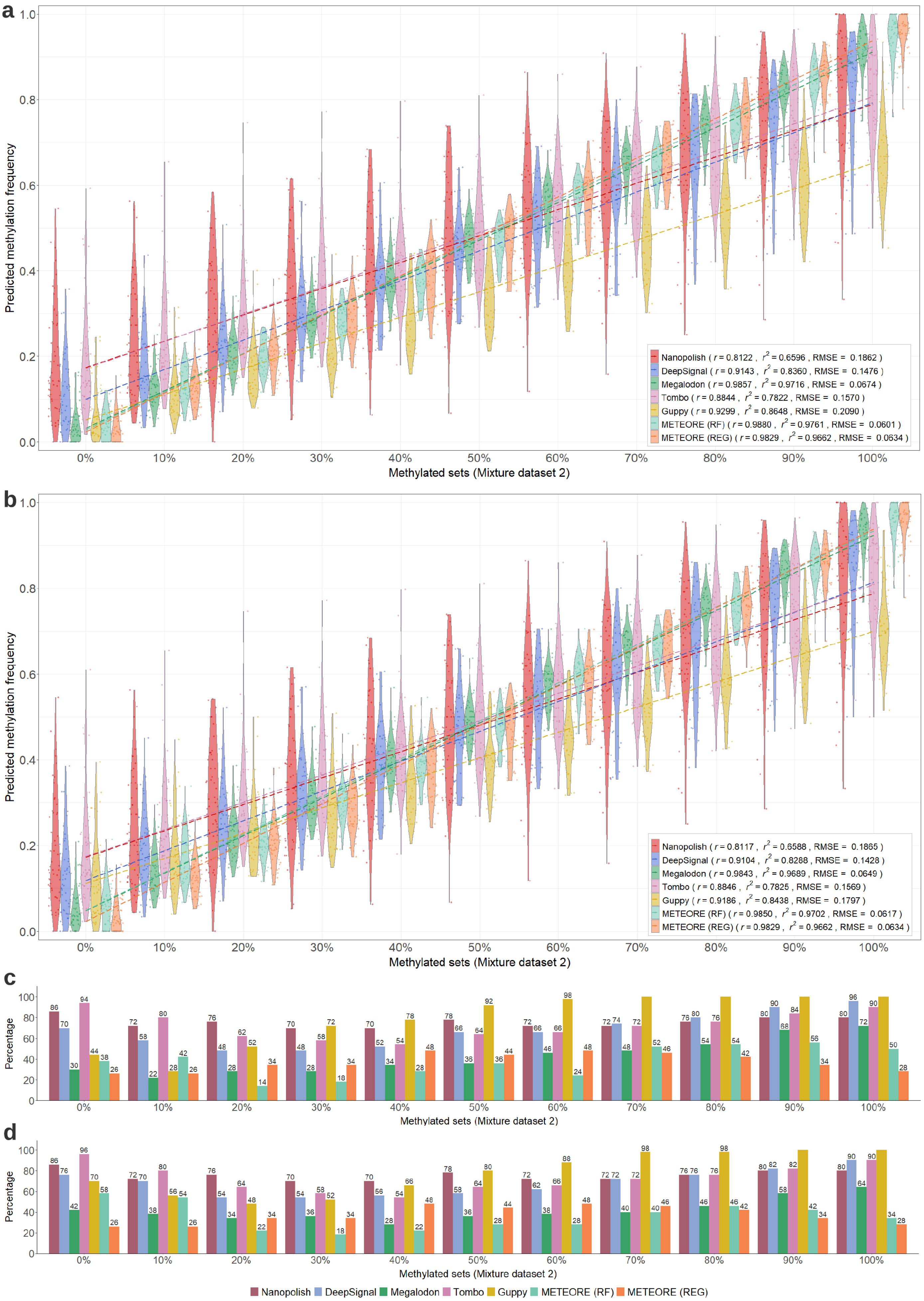
a
b
c
d
Predicted methylation frequency
Methylated sets (Mixture dataset 2)
Percentage
Nanopolish ( r = 0.8122 , r2 = 0.6596 , RMSE = 0.1862 )
DeepSignal ( r = 0.9143 , r2 = 0.8360 , RMSE = 0.1476 )
Megalodon ( r = 0.9857 , r2 = 0.9716 , RMSE = 0.0674 )
Tombo ( r = 0.8844 , r2 = 0.7822 , RMSE = 0.1570 )
Guppy ( r = 0.9299 , r2 = 0.8648 , RMSE = 0.2090 )
METEORE (RF) ( r = 0.9880 , r2 = 0.9761 , RMSE = 0.0601 )
METEORE (REG) ( r = 0.9829 , r2 = 0.9662 , RMSE = 0.0634 )
Nanopolish ( r = 0.8117 , r2 = 0.6588 , RMSE = 0.1865 )
DeepSignal ( r = 0.9104 , r2 = 0.8288 , RMSE = 0.1428 )
Megalodon ( r = 0.9843 , r2 = 0.9689 , RMSE = 0.0649 )
Tombo ( r = 0.8846 , r2 = 0.7825 , RMSE = 0.1569 )
Guppy ( r = 0.9186 , r2 = 0.8438 , RMSE = 0.1797 )
METEORE (RF) ( r = 0.9850 , r2 = 0.9702 , RMSE = 0.0617 )
METEORE (REG) ( r = 0.9829 , r2 = 0.9662 , RMSE = 0.0634 )
Nanopolish DeepSignal Megalodon Tombo Guppy METEORE (RF) METEORE (REG)

**Supplementary Figure 5. Accuracy analysis of five individual tested tools using a single score cutoff. a, b** Violin plots showing the predicted methylation frequencies (y axis) for each control mixture set with a given proportion of methylated reads (x axis) from the mixture dataset 2 for the five tested tools plus METEORE combining Megalodon and DeepSignal using random forest (RF) and regression (REG) models with the single threshold obtained by **(a)** the maximum value of (TPR-FPR) or **(b)** the minimum value of $FPR^2 + (1-TPR)^2$. Score thresholds are given in Supplementary Table 3. The Pearson's correlation (r), coefficient of determination ($r^2$) and the root mean square error (RMSE) are given for each tool. **c, d** Barplot showing the proportion of sites predicted outside a 10% window around the expected methylation proportion for each method with the single threshold obtained by **(c)** the maximum value of (TPR-FPR) or **(d)** the minimum value of $FPR^2 + (1-TPR)^2$. Each predicted site in the m% dataset was classified as "outside" if its predicted percentage methylation was outside the interval [ (m-5)%, (m+5)% ] for intermediate methylation values, or outside the intervals [0,5%] or [95%,100%] for the fully unmethylated or fully methylated sets, respectively. TPR = true positive rate, FPR = false positive rate.

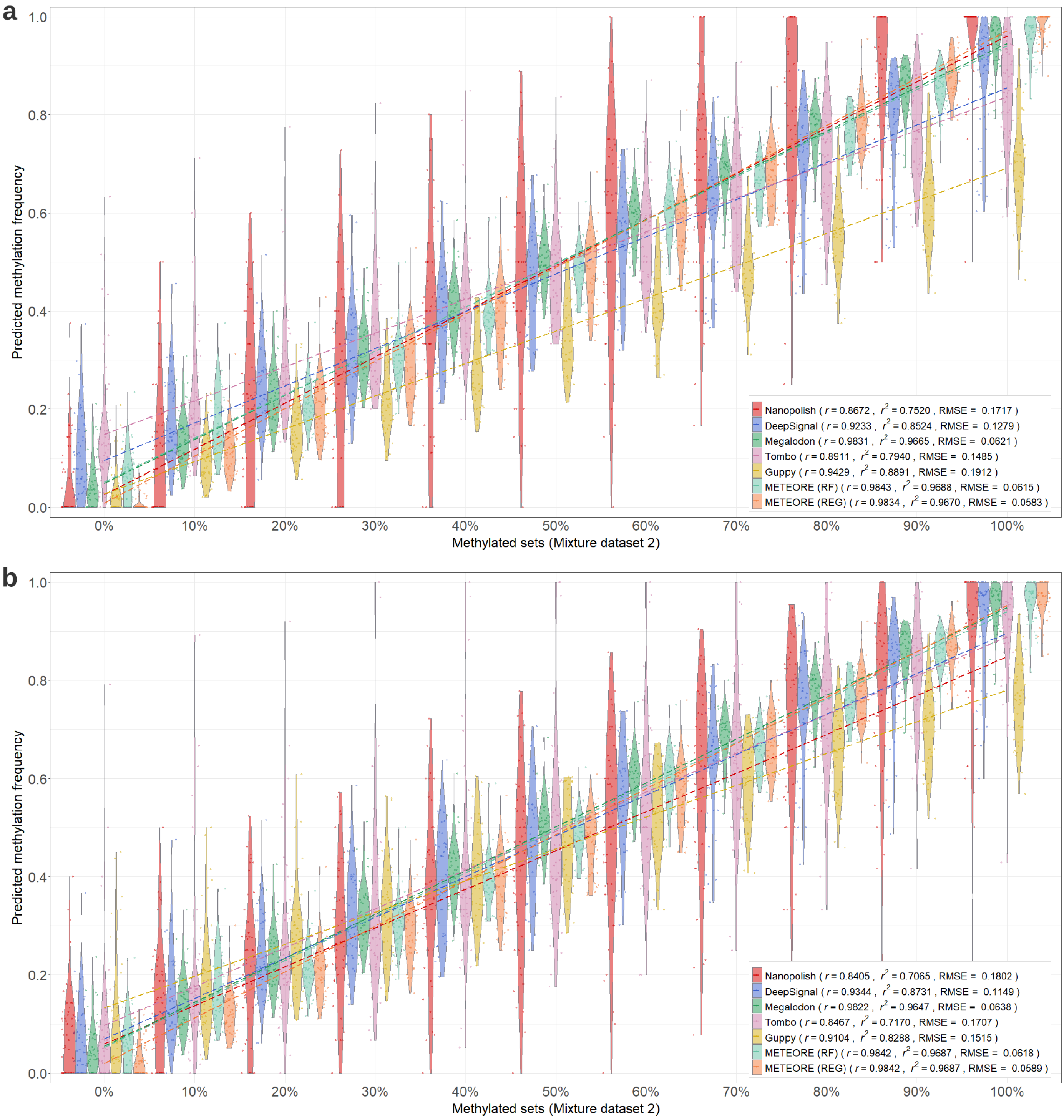
a
b
Predicted methylation frequency
1.0
0.8
0.6
0.4
0.2
0.0
0%
10%
20%
30%
40%
50%
60%
70%
80%
90%
100%
Methylated sets (Mixture dataset 2)
Nanopolish ( r = 0.8672 , r2 = 0.7520 , RMSE = 0.1717 )
DeepSignal ( r = 0.9233 , r2 = 0.8524 , RMSE = 0.1279 )
Megalodon ( r = 0.9831 , r2 = 0.9665 , RMSE = 0.0621 )
Tombo ( r = 0.8911 , r2 = 0.7940 , RMSE = 0.1485 )
Guppy ( r = 0.9429 , r2 = 0.8891 , RMSE = 0.1912 )
METEORE (RF) ( r = 0.9843 , r2 = 0.9688 , RMSE = 0.0615 )
METEORE (REG) ( r = 0.9834 , r2 = 0.9670 , RMSE = 0.0583 )
Nanopolish ( r = 0.8405 , r2 = 0.7065 , RMSE = 0.1802 )
DeepSignal ( r = 0.9344 , r2 = 0.8731 , RMSE = 0.1149 )
Megalodon ( r = 0.9822 , r2 = 0.9647 , RMSE = 0.0638 )
Tombo ( r = 0.8467 , r2 = 0.7170 , RMSE = 0.1707 )
Guppy ( r = 0.9104 , r2 = 0.8288 , RMSE = 0.1515 )
METEORE (RF) ( r = 0.9842 , r2 = 0.9687 , RMSE = 0.0618 )
METEORE (REG) ( r = 0.9842 , r2 = 0.9687 , RMSE = 0.0589 )

c

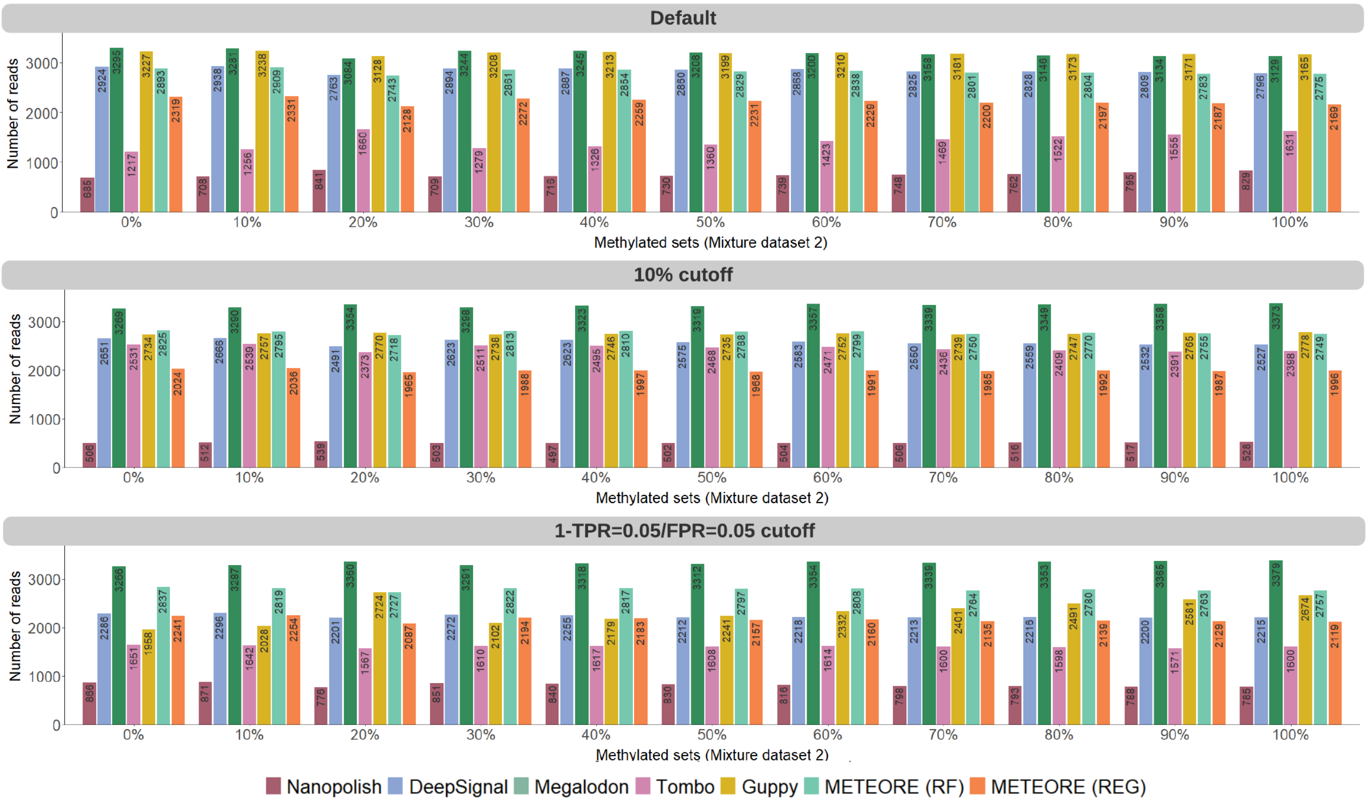


**Supplementary Figure 6. Accuracy analysis using a double cutoff and discarding reads. a** Violin plots showing the predicted methylation frequencies (y axis) for each control mixture set with a given proportion of methylated reads (x axis) from the mixture dataset 2 for the five tested tools plus METEORE combining Megalodon and DeepSignal using random forest (RF) and regression (REG) models, after discarding 10% of the reads with a score closest to the value corresponding to the intersection between FPR and 1-TPR. The Pearson's correlation (r), coefficient of determination ($r^2$) and the root mean square error (RMSE) are given for each tool. **b** Similar plot as **(a)** but considering the scores at which FPR=0.05 and 1-TPR=0.05 and removing all sites in reads with a score between these two values. Cutoffs are given in Supplementary Table 4. **c** Total number of reads reported by each method in different approaches: default setting of each method (top), the use of a double cutoff to remove 20% of the reads (middle) and the use of a double cutoff to remove reads with the scores between the cutoff values at FPR=0.05 and 1-TPR=0.05 (bottom).

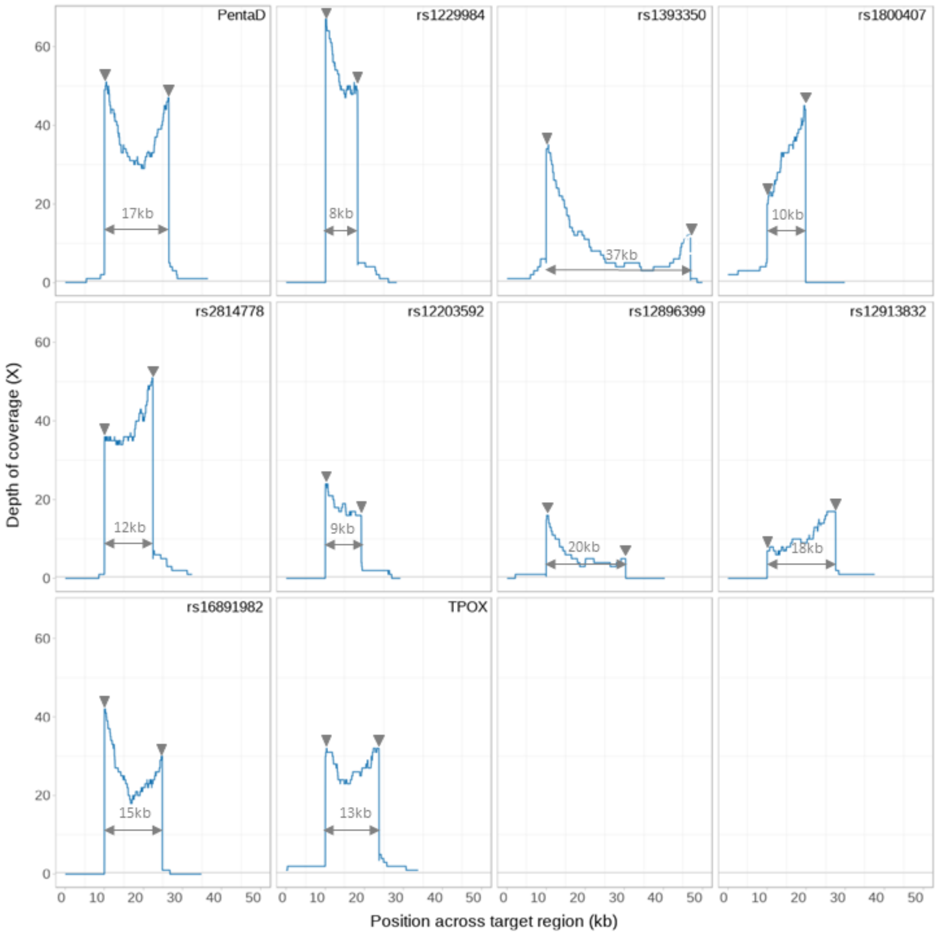


**Supplementary Figure 7. Coverage plots for the 10 regions targeted with the nCATS protocol.** For each of our 10 sequenced regions (Supplementary Table 5), we show the number of reads (y axis) aligning at each position along the region (x axis). The boundaries and length of each region are also indicated. For the coverage, reads mapped in forward and reverse were considered.

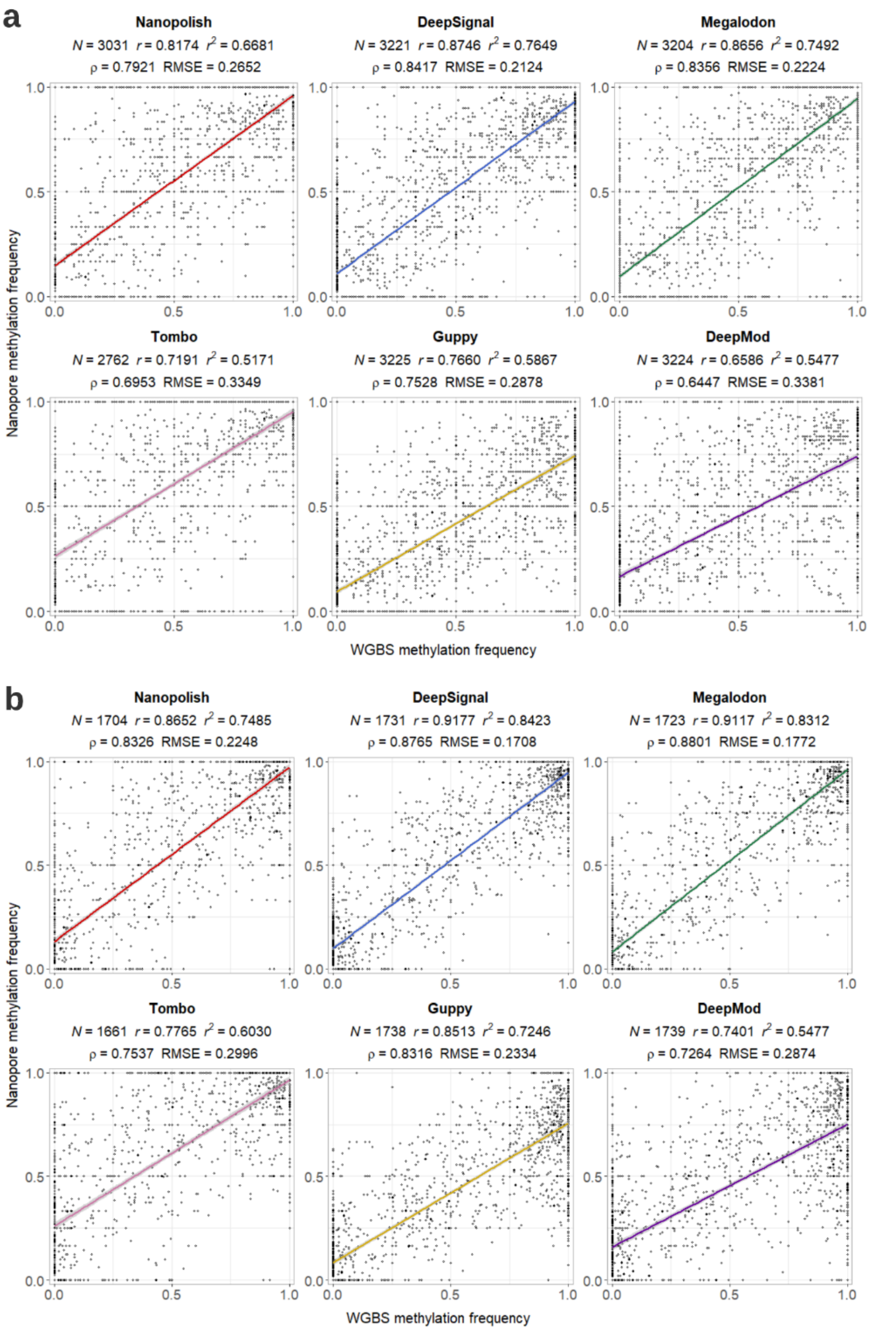


**Supplementary Figure 8. Comparison of the nCATS data with whole genome bisulfite sequencing (WGBS).** For each tool, we show the methylation fraction predicted by each tool (y axis) and the fraction calculated from WGBS (x axis), using either **(a)** individual predictions on both strands or **(b)** combined predictions from both strands. The number of sites (N), the Pearson's correlation (r), coefficient of determination ($r^2$), the Spearman's rank correlation (ρ), and the root mean square error (RMSE) are provided for each tested tool. The plots include the correlation bands.

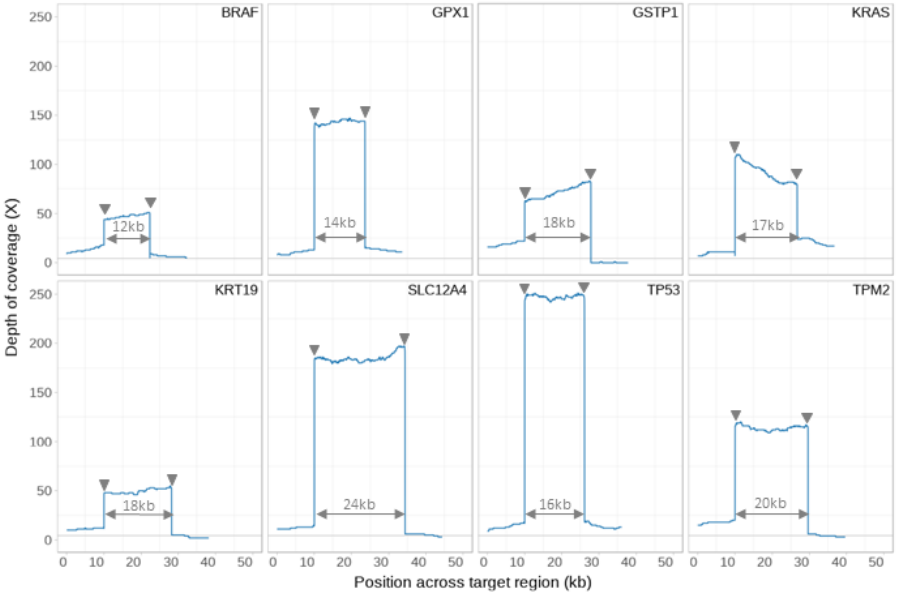


**Supplementary Figure 9. Coverage plots for the regions targeted with the nCATS protocol from Gilpatrick et al. 2020.** For each of the 8 regions tested in Gilpatrick et al. (2020), we show the number of reads (y axis) aligning at each position along the region (x axis). The boundaries and length of each region are also indicated. For the coverage, reads mapped in forward and reverse were considered.

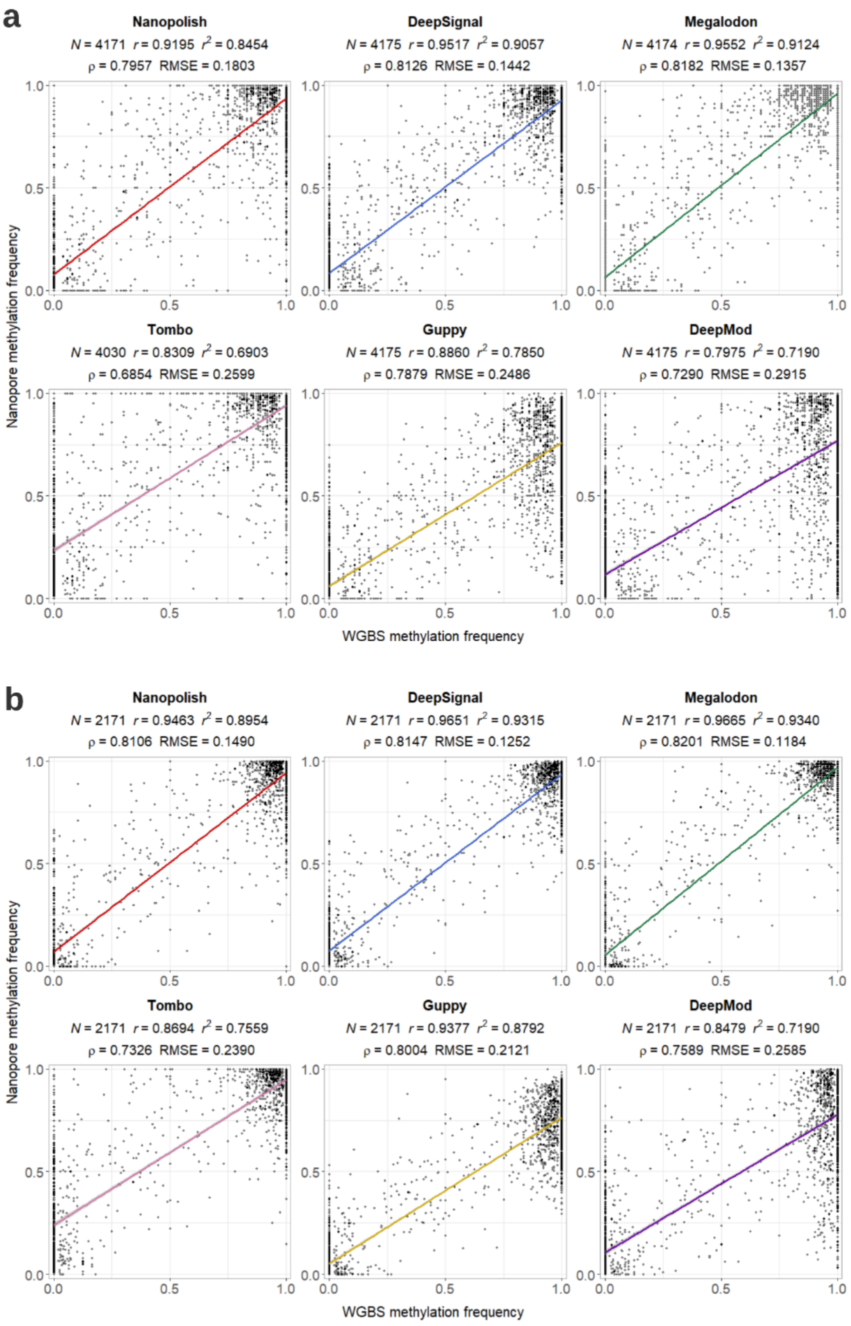


**Supplementary Figure 10. Comparison of the nCATS data from Gilpatrick et al. with whole genome bisulfite sequencing (WGBS) data.** For each tool, we show the methylation fraction predicted by each tool (y axis) and the fraction calculated from WGBS (x axis), using either **(a)** individual predictions on both strands or **(b)** combined predictions from both strands. The number of sites (N), the Pearson's correlation (r), coefficient of determination ($r^2$), the Spearman's rank correlation (ρ), and the root mean square error (RMSE) are provided for each tested tool. The plots include the correlation bands.

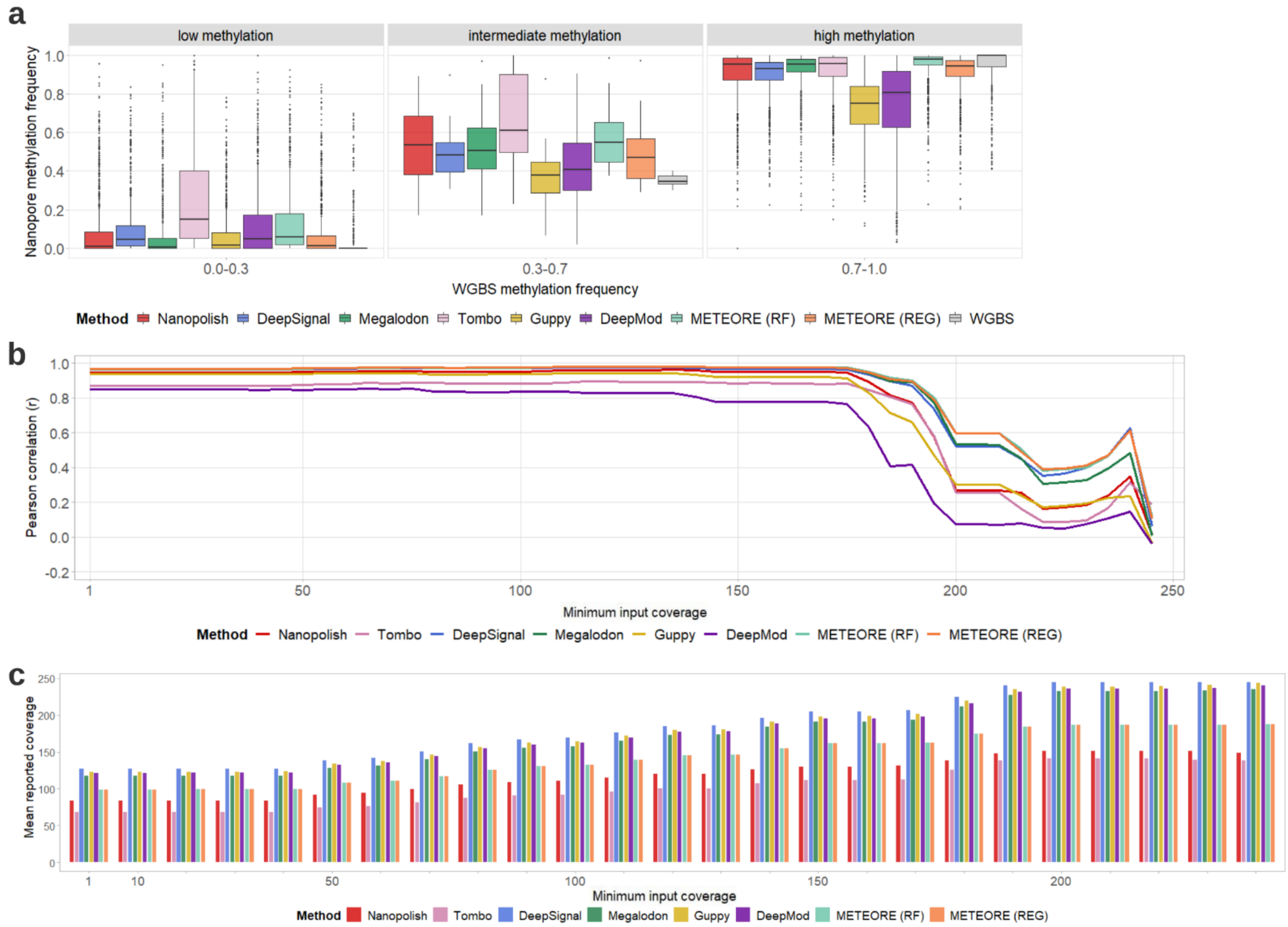


**Supplementary Figure 11. Comparison of CpG methylation frequencies from whole genome bisulfite sequencing (WGBS) Illumina data with Cas9-targeted Nanopore data from Gilpatrick et al. a** Distribution of Nanopore methylation calls across three WGBS methylation bins unmethylated or lowly methylated (0.0-0.3), intermediate methylation (0.3-0.7), and highly or fully methylated (0.7-1.0). We show the seven tested tools: Nanopolish, DeepSignal, Megalodon, Tombo, Guppy, DeepMod, and METEORE. For METEORE, we used the combination of Megalodon and DeepSignal using either a random forest model (RF) or a regression model (REG). **b** Pearson's correlation (r) (y axis) between methylation frequencies calculated from Nanopore by each of the tested tools and WGBS at sites with predictions from both strands combined at each level of minimal input coverage (x axis), i.e., minimum number of Nanopore reads considered per site as reported from the BAM file. **c** Mean reported coverage (y axis), using the coverage reported by each tool for each site, at each value of minimum input coverage in (b) (x axis). METEORE (RF) is the combination of DeepSignal and Megalodon using a random forest (parameters: max_depth=3 and n_estimator=10). METEORE (REG) is the combination of DeepSignal and Megalodon using a regression model.

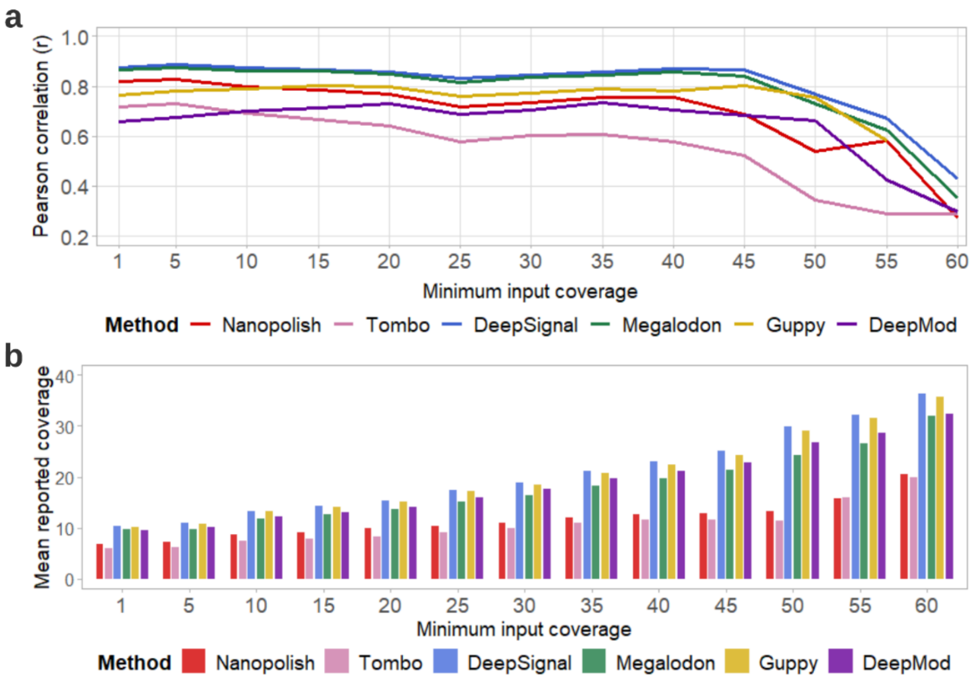


**Supplementary Figure 12. Comparison of CpG methylation frequencies from whole genome bisulfite sequencing (WGBS) Illumina data with Cas9-targeted Nanopore data independently on each strand. a** Pearson's correlation (r) (y axis) between methylation frequencies calculated from Nanopore by each of the tested tools and WGBS at individual sites at each level of minimal input coverage, i.e. minimum number of Nanopore reads considered per site as reported from the BAM file (x axis). **b** Mean reported coverage (using the coverage reported by each tool for each site) considered at each value of minimum input coverage in (a). METEORE is not included since it performs predictions only combining both strands.

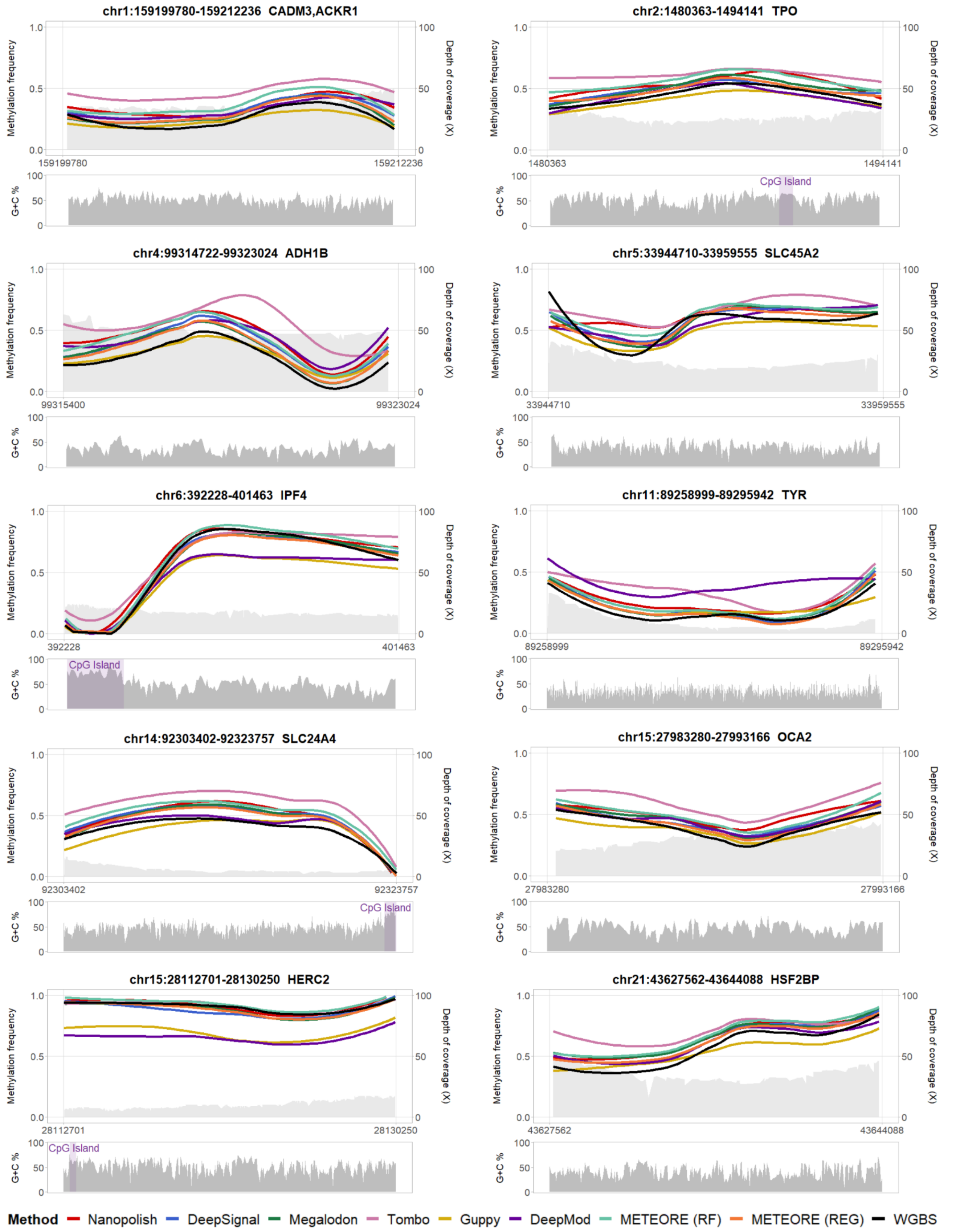


**Supplementary Figure 13. Comparison of CpG methylation frequencies from whole genome bisulfite data (WGBS) and Nanopore across our 10 targeted regions.** Locally Weighted Scatterplot Smoothing (LOESS) smoothing line plots of methylation calls from WGBS Illumina and Nanopore data detected by the eight tested tools. We show METEORE with the combination of Megalodon and DeepSignal using either a random forest model (RF) or a regression model (REG). The plots include the Nanopore coverage, shown as a light grey area. Below, we include the GC-content of the region.

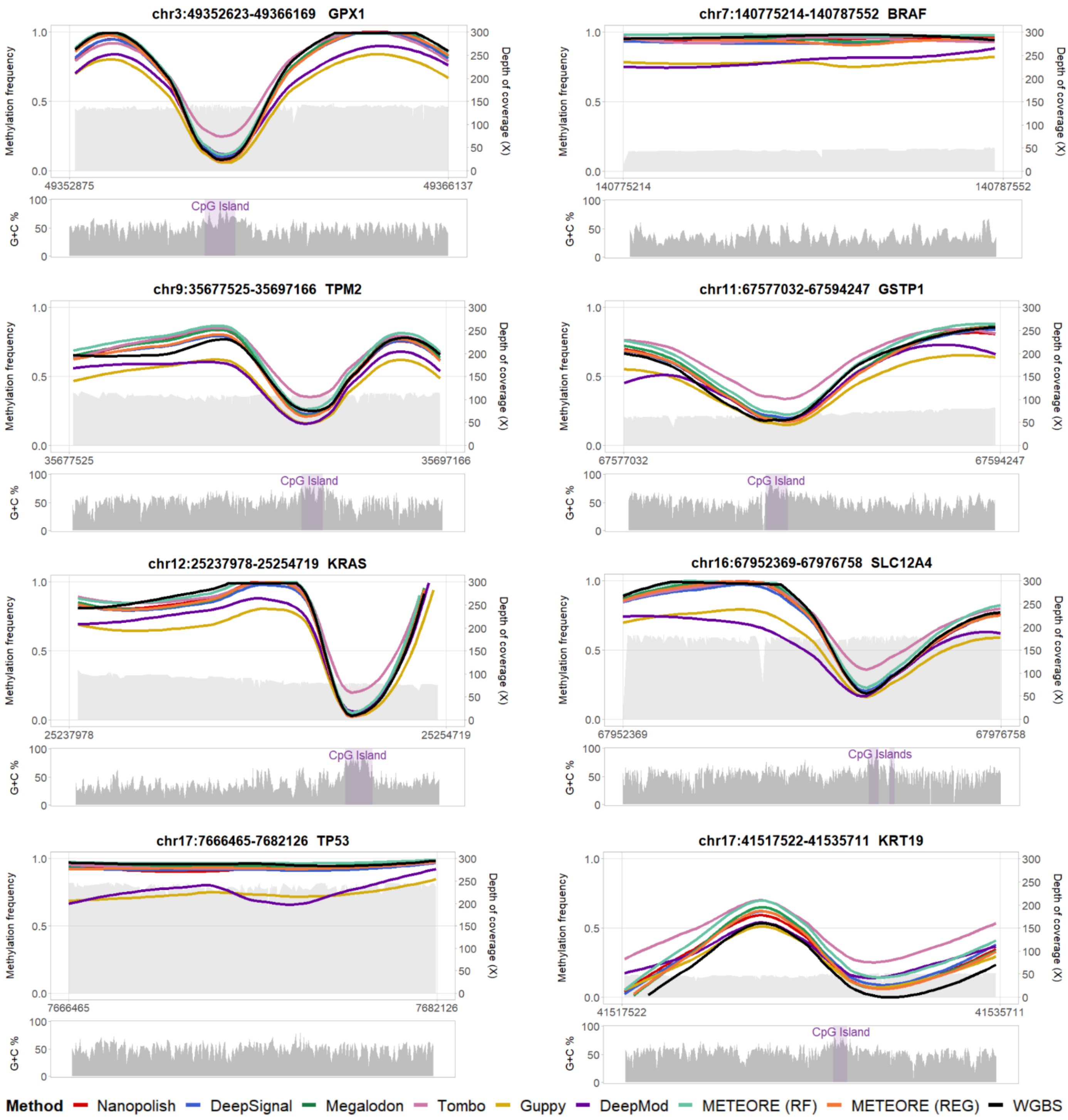


**Supplementary Figure 14. Comparison of CpG methylation frequencies from whole genome bisulfite data (WGBS) and Nanopore across the 8 regions tested in Gilpatrick et al.** Locally Weighted Scatterplot Smoothing (LOESS) smoothing line plots of methylation calls from WGBS Illumina and Nanopore data detected by the eight tested tools. We show METEORE with the combination of Megalodon and DeepSignal using either a random forest model (RF) or a regression model (REG). The plots include the Nanopore coverage, shown as a light grey area. Below, we include the GC-content of the region.

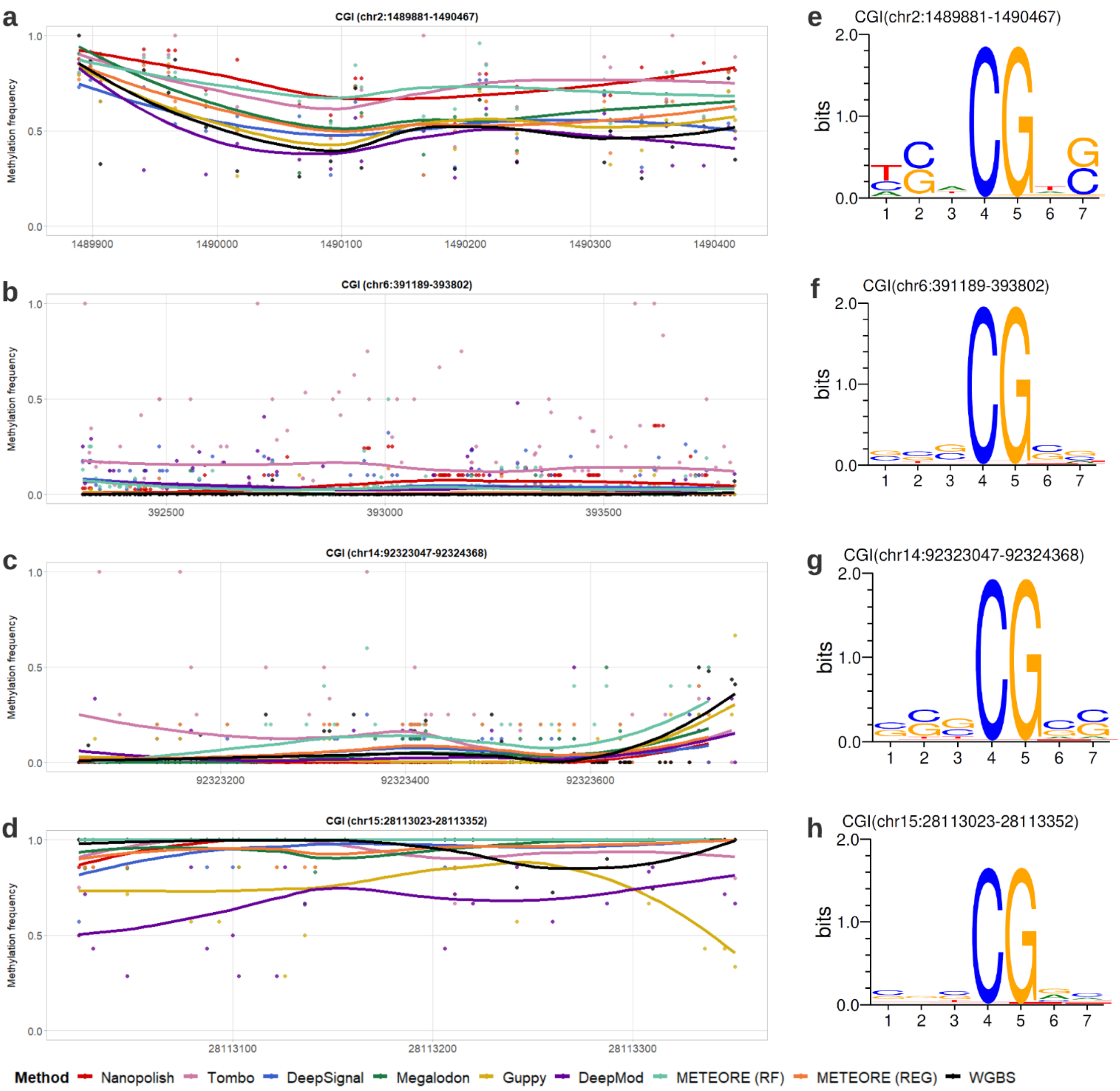


**Supplementary Figure 15. Zoom in on the CpG Islands (CGIs) for the comparison of CpG methylation predictions from Nanopore with whole genome bisulfite sequencing (WGBS). a-d** show a zoom in of the LOESS smoothing line plots of methylation frequency (y axis) with individual methylation calls (points) for nine different methods for the CGIs in Figure 5. **e-h** show the sequence logos showing information content in bits of the motifs (7-mers) at the CpG sites in the same CGIs shown in (a-d).

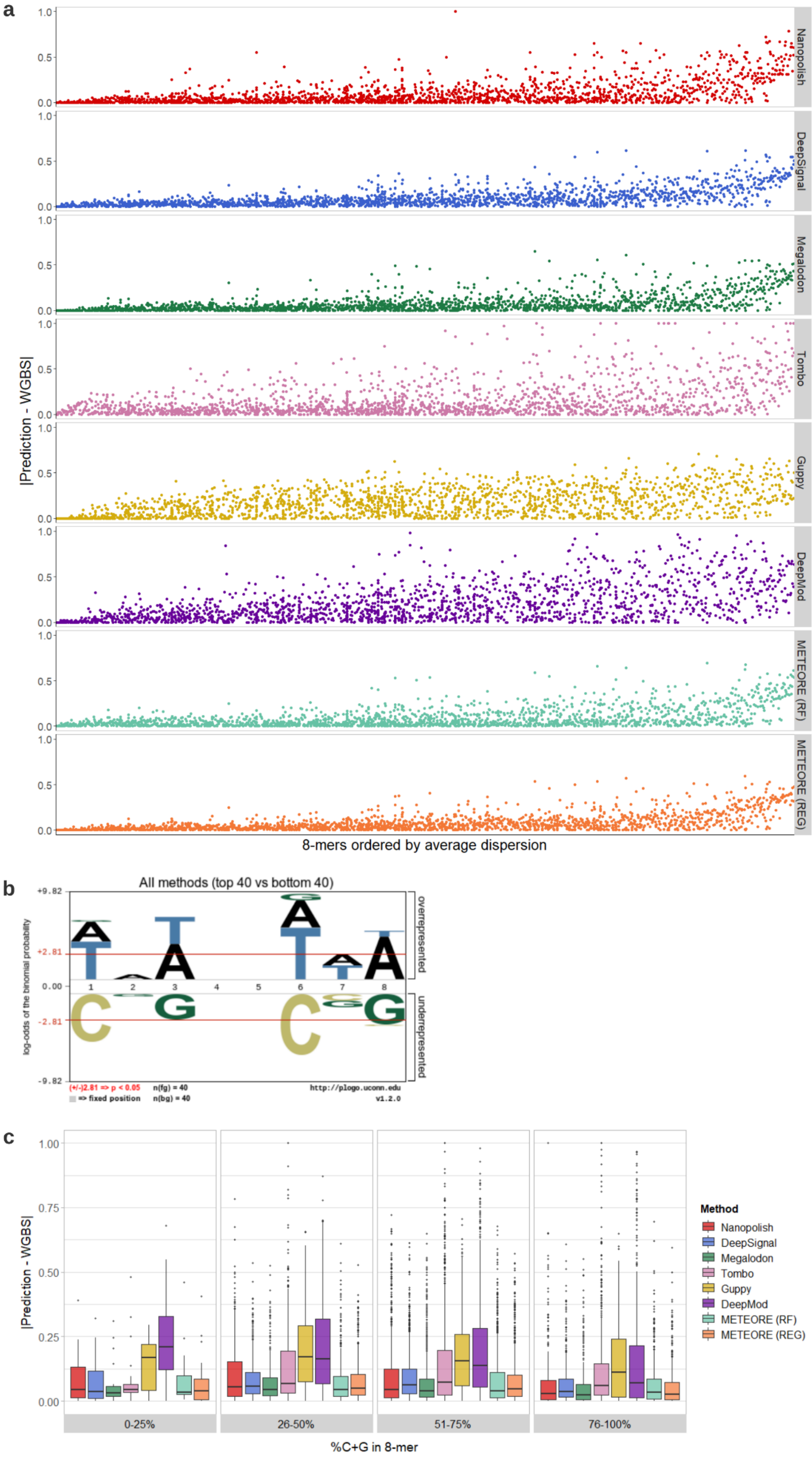


**Supplementary Figure 16. Sequence context analysis.** Absolute difference between methylation frequencies from Nanopore-based methylation detection tools and whole genome bisulfite sequencing (WGBS) data for all 8-mers with a CpG in the middle (NNNCGNNN) (x axis) ordered from left to right in ascending order according to the average value of the absolute difference (y axis) across all tools. **b** Sequence logo generated by the plogo Web tool for the top/bottom 40 8-mers from the ranking in (a). The bottom 40 8-mers were considered the foreground (fg) dataset, as shown in the upper panel, whereas the top 40 8-mers where considered the background (bg) dataset, as shown in the lower panel. Residues are stacked according to the statistical significance, with the most significant residues positioned closest to the x axis. The significance level (α) and the number of foreground and background sequences used, i.e., n(fg) and n(bg) values, are given at the bottom of each logo. The red horizontal lines correspond to $\alpha = 0.05$. **c** Same absolute difference of methylation frequencies in 8-mers from (a) (y axis) but stratified by the percentage of C and G residues in the 8-mers, separated by tool.

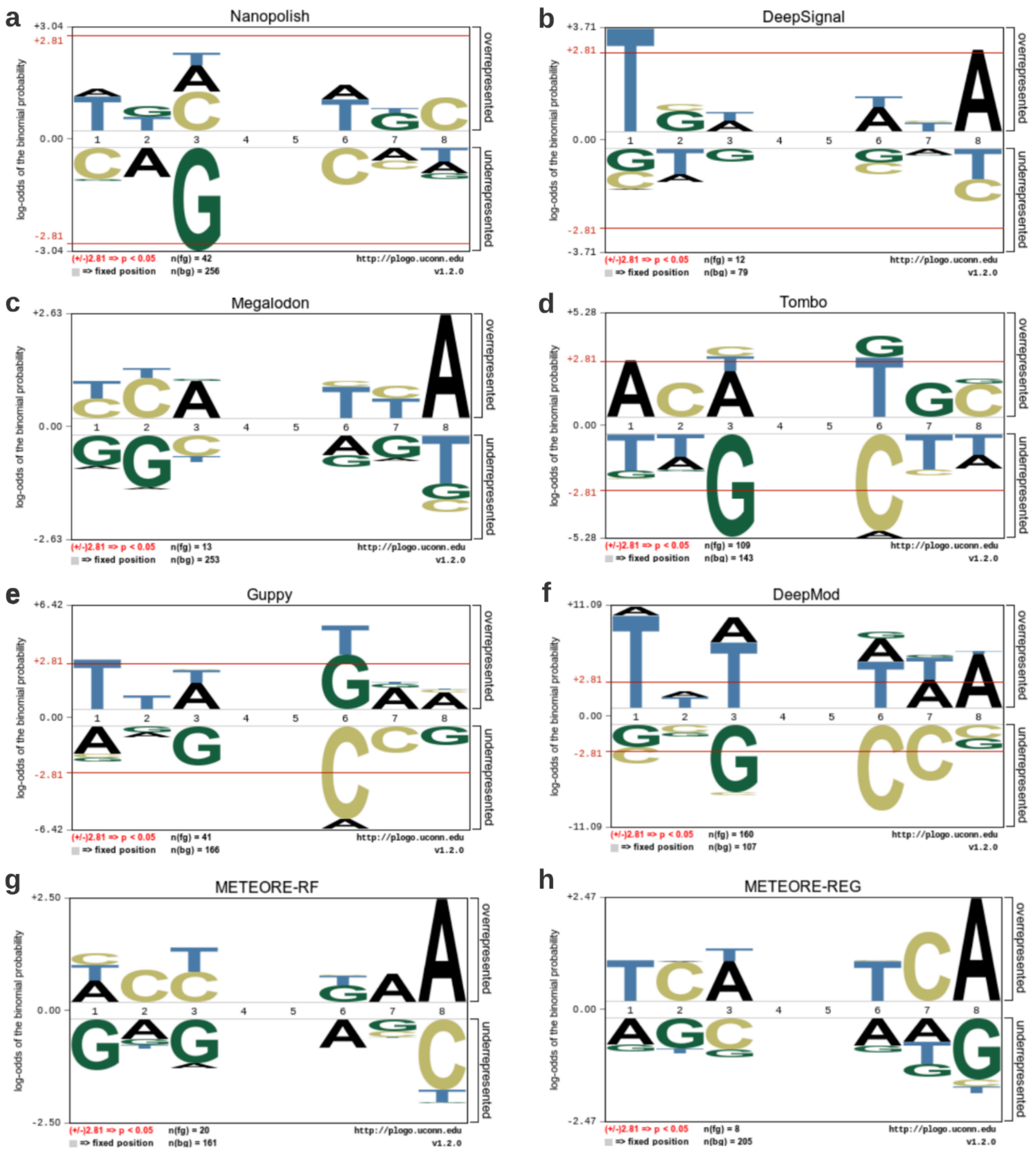


**Supplementary Figure 17. Tests of the sequences associated to sites of high and low discrepancy with whole genome bisulfite sequencing (WGBS).** For each method, CG-containing 8-mers (NNNCGNNN) were labelled as "bad" (foreground (fg) dataset, as shown in the upper panel of each plot) if the discrepancy with WGBS (in absolute value) was >0.5, or as "good" (background (bg) dataset, as shown in the lower panel of each plot) if the discrepancy with WGBS was exactly 0. Residues are stacked according to the statistical significance, with the most significant residues positioned closest to the x axis. The significance level (α) and the number of foreground and background sequences used, i.e., n(fg) and n(bg) values, are given at the bottom of each logo. The red horizontal lines correspond to α = 0.05.

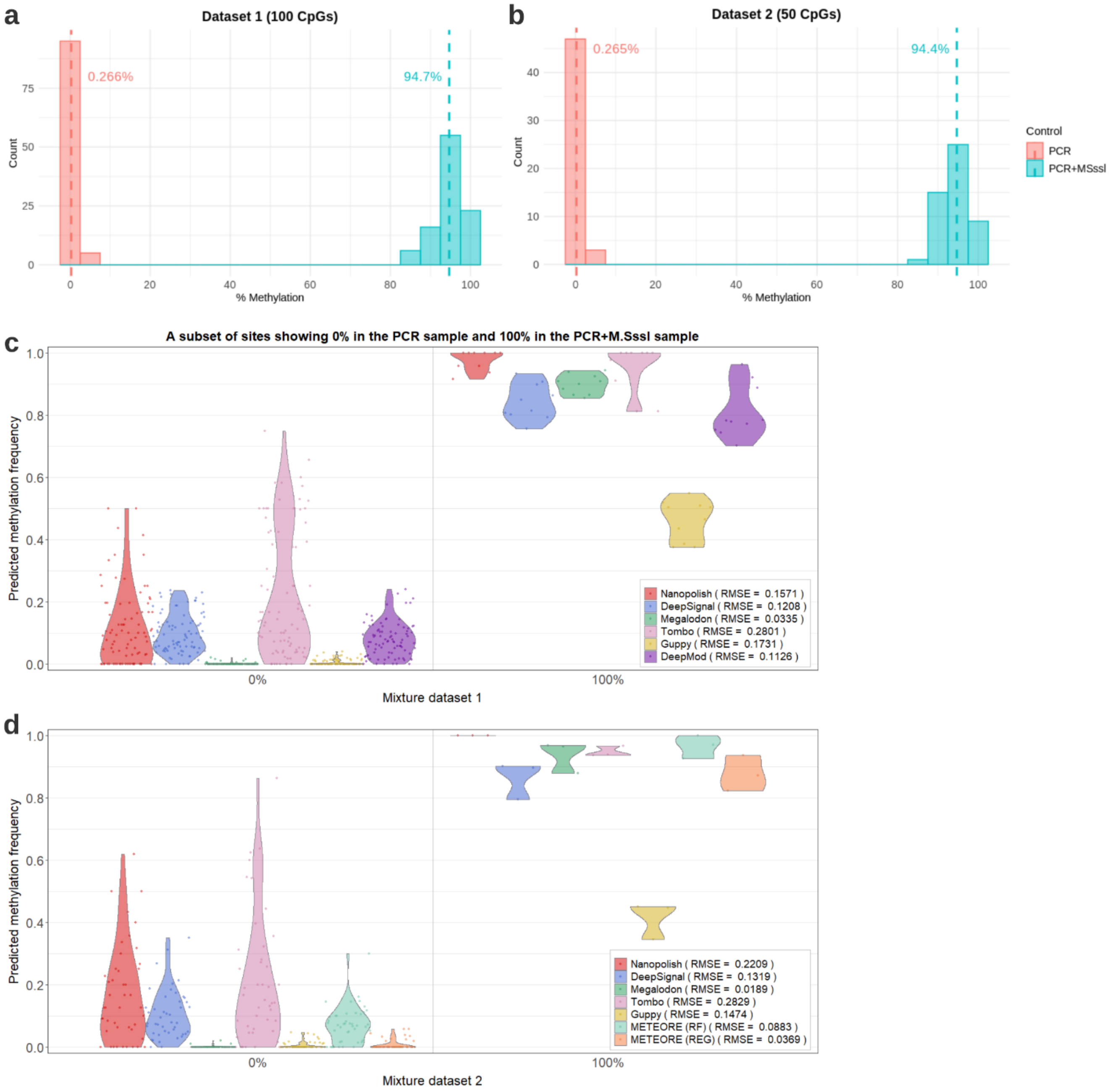


**Supplementary Figure 18. Confirmation of the methylation levels of control samples using whole genome bisulfite sequencing (WGBS) data. (a)** Distribution plot showing the methylation percentage for the 100 target sites in the mixture dataset 1 in the positive (PCR+M.SssI) (mean methylation = 94.7%) and negative (PCR) (mean methylation = 0.266%) controls. **(b)** Distribution plot showing methylation percentage for the 50 target sites in the mixture dataset 2 in the positive (mean = 94.4%) and negative (mean = 0.265%) controls. **(c)** Violin plots showing the distribution of predicted methylation frequencies (y axis) in the sites from dataset 1 that in (a) showed exactly 0% methylation in the negative control and exactly 100% methylation in the positive control, using WGBS. The root mean square error (RMSE) is given for each tool. **(d)** Violin plots showing the distribution of predicted methylation frequencies (y axis) in the sites from the mixture dataset 2 that in (b) showed exactly 0% methylation in the negative control and exactly 100% methylation in the positive control, using WGBS.

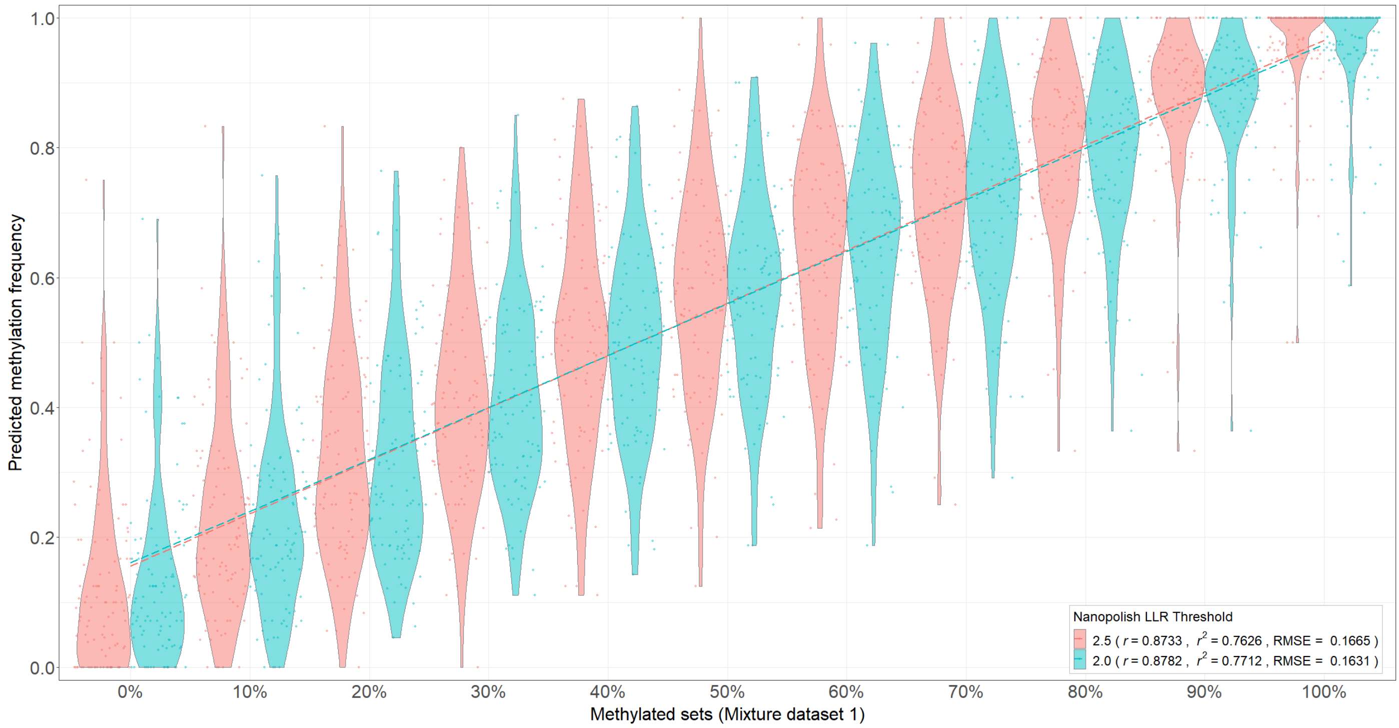


**Supplementary Figure 19. Comparison of methylation frequencies using log-likelihood ratio (LLR) thresholds of 2.5 and 2.0 for Nanopolish.** Two different LLR thresholds were used to make a per-site methylation call on control mixture dataset 1.

a

| Set name | % methylated | Total reads | Unmethylated reads (PCR) | Methylated reads (PCR+M.SssI) |
|---|---|---|---|---|
| m0 | 0 | 2390 | 2390 | 0 |
| m10 | 10 | 2437 | 2182 | 255 |
| m20 | 20 | 2431 | 1946 | 485 |
| m30 | 30 | 2434 | 1714 | 720 |
| m40 | 40 | 2410 | 1458 | 952 |
| m50 | 50 | 2432 | 1231 | 1201 |
| m60 | 60 | 2414 | 981 | 1433 |
| m70 | 70 | 2420 | 739 | 1681 |
| m80 | 80 | 2410 | 498 | 1912 |
| m90 | 90 | 2399 | 256 | 2143 |
| m100 | 100 | 2225 | 0 | 2225 |

b

| Set name | % methylated | Total reads | Unmethylated reads (PCR) | Methylated reads (PCR+M.SssI) |
|---|---|---|---|---|
| m0 | 0 | 3420 | 3420 | 0 |
| m10 | 10 | 3426 | 3081 | 345 |
| m20 | 20 | 3413 | 2745 | 668 |
| m30 | 30 | 3423 | 2410 | 1013 |
| m40 | 40 | 3415 | 2068 | 1347 |
| m50 | 50 | 3434 | 1736 | 1698 |
| m60 | 60 | 3403 | 1377 | 2026 |
| m70 | 70 | 3406 | 1039 | 2367 |
| m80 | 80 | 3398 | 690 | 2708 |
| m90 | 90 | 3396 | 354 | 3042 |
| m100 | 100 | 3383 | 0 | 3383 |

**Supplementary Table 1. Methylation control mixtures.** We describe the mixture dataset 1 **(a)** and 2 **(b)** used for the benchmarking of different methylation proportions built from fully unmethylated and fully methylated reads.

| Unmethylated if freq < 0.1, methylated if freq > 0.9 | | | | | |
|---|---|---|---|---|---|
| | Accuracy | Specificity | Precision | Recall | Error rate |
| **Nanopolish** | 0.675 | 0.490 | 0.628 | 0.860 | 0.325 |
| **DeepSignal** | 0.410 | 0.590 | 0.359 | 0.230 | 0.590 |
| **Megalodon** | 0.825 | 1.000 | 1.000 | 0.650 | 0.175 |
| **Tombo** | 0.588 | 0.384 | 0.564 | 0.790 | 0.412 |
| **Guppy** | 0.500 | 1.000 | NA | 0.000 | 0.500 |
| **DeepMod** | 0.495 | 0.680 | 0.492 | 0.310 | 0.505 |

| Unmethylated if freq < 0.2, methylated if freq > 0.8 | | | | | |
|---|---|---|---|---|---|
| | Accuracy | Specificity | Precision | Recall | Error rate |
| **Nanopolish** | 0.860 | 0.800 | 0.821 | 0.920 | 0.140 |
| **DeepSignal** | 0.805 | 0.910 | 0.886 | 0.700 | 0.195 |
| **Megalodon** | 0.970 | 1.000 | 1.000 | 0.940 | 0.030 |
| **Tombo** | 0.794 | 0.616 | 0.719 | 0.970 | 0.206 |
| **Guppy** | 0.500 | 1.000 | NA | 0.000 | 0.500 |
| **DeepMod** | 0.790 | 0.970 | 0.953 | 0.610 | 0.210 |

| Unmethylated if freq < 0.3, methylated if freq > 0.7 | | | | | |
|---|---|---|---|---|---|
| | Accuracy | Specificity | Precision | Recall | Error rate |
| **Nanopolish** | 0.935 | 0.900 | 0.907 | 0.970 | 0.065 |
| **DeepSignal** | 0.930 | 1.000 | 1.000 | 0.860 | 0.070 |
| **Megalodon** | 0.980 | 1.000 | 1.000 | 0.960 | 0.020 |
| **Tombo** | 0.839 | 0.697 | 0.766 | 0.980 | 0.161 |
| **Guppy** | 0.505 | 1.000 | 1.000 | 0.010 | 0.495 |
| **DeepMod** | 0.935 | 1.000 | 1.000 | 0.870 | 0.065 |

| Unmethylated if freq < 0.4, methylated if freq > 0.6 | | | | | |
|---|---|---|---|---|---|
| | Accuracy | Specificity | Precision | Recall | Error rate |
| **Nanopolish** | 0.955 | 0.930 | 0.933 | 0.980 | 0.045 |
| **DeepSignal** | 0.975 | 1.000 | 1.000 | 0.950 | 0.025 |
| **Megalodon** | 1.000 | 1.000 | 1.000 | 1.000 | 0.000 |
| **Tombo** | 0.884 | 0.778 | 0.818 | 0.990 | 0.116 |
| **Guppy** | 0.600 | 1.000 | 1.000 | 0.200 | 0.400 |
| **DeepMod** | 0.975 | 1.000 | 1.000 | 0.950 | 0.025 |

| Unmethylated if freq < 0.5, methylated if freq > 0.5 | | | | | |
|---|---|---|---|---|---|
| | Accuracy | Specificity | Precision | Recall | Error rate |
| **Nanopolish** | 0.975 | 0.970 | 0.970 | 0.980 | 0.025 |
| **DeepSignal** | 0.990 | 1.000 | 1.000 | 0.980 | 0.010 |
| **Megalodon** | 1.000 | 1.000 | 1.000 | 1.000 | 0.000 |
| **Tombo** | 0.915 | 0.838 | 0.861 | 0.990 | 0.085 |
| **Guppy** | 0.720 | 1.000 | 1.000 | 0.440 | 0.280 |
| **DeepMod** | 0.980 | 1.000 | 1.000 | 0.960 | 0.020 |

**Supplementary Table 2. Per-site performance.** The table shows the accuracies in fully methylated or fully unmethylated CpG sites for the six tested tools using two different methylation frequency thresholds to classify methylated and unmethylated sites. For each pair of thresholds (a,b), we defined a site to be unmethylated if the predicted methylation frequency was <a, and methylated if the predicted methylation frequency was >b, for (a,b) = (0.1,0.9), (0.2,0.8), (0.3,0.7), (0.4,0.6), and (0.5,0.5).

| | Nanopolish | DeepSignal | Megalodon | Tombo | Guppy | METEORE (RF) | METEORE (REG) |
|---|---|---|---|---|---|---|---|
| Maximum of (TPR –FPR) | Cutoff= 1.03<br>TPR =0.87<br>FPR=0.15 | Cutoff= -0.05<br>TPR =0.86<br>FPR=0.10 | Cutoff= -0.98<br>TPR =0.91<br>FPR=0.04 | Cutoff= -0.13<br>TPR =0.83<br>FPR=0.18 | Cutoff= -1.35<br>TPR =0.69<br>FPR=0.05 | Cutoff= 0.62<br>TPR =0.94<br>FPR=0.03 | Cutoff= 0.33<br>TPR =0.95<br>FPR=0.03 |
| Minimum of $(FPR\text{-}0)^2 + (TPR\text{-}1)^2$ | Cutoff= 1.04<br>TPR=0.87<br>FPR=0.15 | Cutoff= -0.19<br>TPR =0.87<br>FPR=0.11 | Cutoff= -1.36<br>TPR =0.92<br>FPR=0.05 | Cutoff= -0.11<br>TPR =0.82<br>FPR=0.18 | Cutoff= -1.70<br>TPR =0.73<br>FPR=0.12 | Cutoff= 0.55<br>TPR =0.95<br>FPR=0.04 | Cutoff= 0.33<br>TPR =0.96<br>FPR=0.04 |

**Supplementary Table 3. Single score cutoffs.** Cutoffs obtained by maximising the value of TPR-FPR (first row) or by minimizing the value of $(FPR\text{-}0)^2 + (TPR\text{-}1)^2$ (second row). In both optimization we used all reads from the mixture dataset 1. These cutoffs were applied to the per-read data generated by each tool. For all these tools except for Tombo, if a read with a score above the cutoff, we consider it as methylated, and unmethylated for the scores below the cutoff. For Tombo, a read is considered methylated if its score is below the cutoff, and unmethylated for a score above the cutoff. All values were rounded up to 2 decimal places. For METEORE REG, both strategies led to exactly the same cutoffs after rounding up.

| | Nanopolish | DeepSignal | Megalodon | Tombo | Guppy | METEORE (RF) | METEORE (REG) |
|---|---|---|---|---|---|---|---|
| Remove **10%** of reads around the cross point of FPR and 1-TPR curves | (-3.58 ,5.80) | (-0.60, 0.09) | (-2.07, -1.20) | (-0.54,0.34) | (-1.93, -0.91) | (0.47,0.56) | (0.20,0.46) |
| Remove the reads that fall between the score at **1-TPR=0.05** and **FPR=0.05** | (-0.65, 3.53) | (-1.07, 0.60) | (-2.14, -1.29) | (-1.82, 1.66) | (-2.41, -1.31) | (0.48,0.55) | (0.29, 0.34) |

**Supplementary Table 4. Double score cutoffs.** Cutoffs obtained by removing 10% of reads around the intersection point of the FPR curve and 1-TPR curve (first row) or removing the cases that fall between the score at 1- TPR = 0.05 and FPR = 0.05 (second row). In the first optimization we used all reads from the mixture dataset 1. In the second optimization we used the fully methylated and fully unmethylated sets from mixture dataset 1. For each double cutoff (a,b), all sites in reads with score < a are considered unmethylated, with score > b are considered methylated, and all cases between these values are discarded. For Tombo the score scale has the opposite orientation, i.e., a read is considered methylated if its score is < a, and unmethylated for a score > b. METEORE (RF) is the combination of DeepSignal and Megalodon using a random forest (parameters: max_depth=3 and n_estimator=10). METEORE (REG) is the combination of DeepSignal and Megalodon using a regression model. In the REG model, filtering is symmetric by the ranking of scores about the tipping point, i.e., 5% of reads with scores lower than the tipping point and 5% of reads with scores higher than the tipping point.

| Chromosome | Start position | End position | Size (nt) | Target locus | Type of variants | Associated gene(s) |
|---|---|---|---|---|---|---|
| chr1 | 159199780 | 159212236 | 12456 | rs2814778 | aiSNP | CADM3, ACKR1 |
| chr2 | 1480363 | 1494141 | 13778 | TPOX | STR | TPO |
| chr4 | 99314722 | 99323024 | 8302 | rs1229984 | aiSNP | ADH1B |
| chr5 | 33944710 | 33959555 | 14845 | rs16891982 | piSNP | SLC45A2 |
| chr6 | 392228 | 401463 | 9235 | rs12203592 | piSNP | IPF4 |
| chr11 | 89258999 | 89295942 | 36943 | rs1393350 | piSNP | TYR |
| chr14 | 92303402 | 92323757 | 20355 | rs12896399 | piSNP | SLC24A4 |
| chr15 | 27983280 | 27993166 | 9886 | rs1800407 | piSNP | OCA2 |
| chr15 | 28112701 | 28130250 | 17549 | rs12913832 | piSNP | HERC2 |
| chr21 | 43627562 | 43644088 | 16526 | PentaD | STR | HSF2BP |

**Supplementary Table 5. Ten forensically relevant regions used for the nCATS protocol.** The table provides the coordinates (GRCh38) of the ten regions used to sequence native DNA with the nCATS protocol. The table also indicates whether the region contains an ancestry-informative SNP (aiSNP), a phenotypic-informative SNP (piSNP), or a short tandem repeat (STR).

| Target | Guide RNA sequence | PAM | Cleaved site |
|---|---|---|---|
| rs12913832 | CTTGTTCTCAATCCAACGAG | CGG | chr15:28112701(+) |
| | GATCAGATGACCATGTTCGA | AGG | chr15:28130250(-) |
| rs1800407 | GTAGAGCTCTAACTAAGTGG | AGG | chr15:27983280(+) |
| | TATCCAATCCTGCTGACCAG | TGG | chr15:27993166(-) |
| rs12896399 | GCTGGAACGCCCCATCAACA | CGG | chr14:92303402(+) |
| | GAGTGCAATCAGTGGCCGAG | CGG | chr14:92323757(-) |
| rs16891982 | TGTGATCACCACGACGACAA | CGG | chr5:33944710(+) |
| | GAGTGCAACGAGGAACTAAG | AGG | chr5:33959555(-) |
| rs1393350 | TCCTTGCTGCACGAATCAGT | GGG | ch11:89258999(+) |
| | GCTGGATGTGTTATAGACGC | TGG | chr11:89295942(-) |
| rs12203592 | TAAGGGGCCCAAGCTCACGG | CGG | chr6:392228(+) |
| | ACGTGGTCAGCTCCTTCACG | AGG | chr6:401463(-) |
| TPOX | CGTATTTGAAAGATCCACGG | TGG | chr2:1480363(+) |
| | CTTACGTAAGAGTTGAATGG | TGG | chr2:1494141(-) |
| Penta D | CGGTACCTATCCCAGAACTA | TGG | chr21:43627562(+) |
| | TAACACGTAGATCATTCACT | TGG | chr21:43644088(-) |
| rs2814778 | CCTACCACGCCATCATCGGT | GGG | chr1:159199780(+) |
| | GCAATTGTCTTTCAGTGCGT | TGG | chr1:159212236(-) |
| rs1229984 | ACCATCTGCTAACACGTATG | AGG | chr4:99314772(+) |
| | GCGTTAACATATCTCCACAA | GGG | chr4:99323024(-) |

**Supplementary Table 6. Guide RNA (gRNA) panel used for the nCATs protocol.** The table describe the ten pairs of gRNAs used to target the ten regions from Supplementary Table 3. To enrich for each target region, two gRNAs were used to make a cut on each side, one upstream of the region of interest targeting the positive strand, and the other one downstream targeting the negative strand.

| | *N* | *r* | $r^2$ | *ρ* | *RMSE* |
|---|---|---|---|---|---|
| Nanopolish | 2171 | 0.9463 | 0.8954 | 0.8106 | 0.1490 |
| DeepSignal | 2171 | 0.9651 | 0.9315 | 0.8147 | 0.1252 |
| Megalodon | 2171 | 0.9665 | 0.9340 | 0.8201 | 0.1184 |
| Tombo | 2171 | 0.8694 | 0.7559 | 0.7326 | 0.2390 |
| Guppy | 2171 | 0.9377 | 0.8792 | 0.8004 | 0.2121 |
| DeepMod | 2171 | 0.8479 | 0.7190 | 0.7589 | 0.2585 |
| METEORE (RF) | 2171 | 0.9641 | 0.9294 | 0.8259 | 0.1307 |
| METEORE (REG) | 2171 | 0.9689 | 0.9387 | 0.8253 | 0.1155 |

**Supplementary Table 7. Comparison of CpG methylation frequencies from whole genome bisulfite sequencing (WGBS) Illumina data with Cas9-targeted Nanopore data from Gilpatrick et al. 2020.** For each tool we provide the number of sites (N), the Pearson's correlation (r), coefficient of determination ($r^2$), the Spearman's rank correlation (ρ), and the root mean square error (RMSE) for the comparison of the percentage methylation predicted from Nanopore with the percentage methylation calculated from WGBS data. We show the results for five tested tools and METEORE combining DeepSignal and Megalodon using a random forest (RF) (parameters: max_depth=3 and n_estimator=10) or a regression (REG) model.

| | 5x | 10x | 20x | 50x |
|---|---|---|---|---|
| Nanopolish | 0.7307 | 0.7719 | 0.7828 | 0.7905 |
| DeepSignal | 0.7851 | 0.8149 | 0.8206 | 0.8294 |
| Megalodon | 0.7999 | 0.8226 | 0.8292 | 0.8371 |
| Guppy | 0.6757 | 0.7161 | 0.7432 | 0.7545 |
| Tombo | 0.6281 | 0.6655 | 0.6892 | 0.7028 |

**Supplementary Table 8. Pearson correlations (r) of methylation frequencies obtained by different Nanopore methylation tools and whole genome bisulfite sequencing (WGBS) at different coverage levels.** We subsampled 5x, 10x, 20x, 50x read coverage for 1487 CpG sites, using Cas9-targeted Nanopore sequencing data from Gilpatrick et al. 2020.

| | *N* | *r* | *$r^2$* | *ρ* | *RMSE* |
|---|---|---|---|---|---|
| Nanopolish | 1724 | 0.8648 | 0.7478 | 0.8362 | 0.2171 |
| DeepSignal | 1731 | 0.9196 | 0.8456 | 0.8785 | 0.1693 |
| Megalodon | 1723 | 0.9040 | 0.8172 | 0.8753 | 0.1938 |
| Tombo | 1734 | 0.8037 | 0.6460 | 0.7871 | 0.2551 |
| Guppy | 1738 | 0.7706 | 0.5938 | 0.7741 | 0.2978 |
| METEORE (RF) | 1723 | 0.9217 | 0.8496 | 0.8878 | 0.1736 |
| METEORE (REG) | 1723 | 0.9164 | 0.8397 | 0.8871 | 0.1829 |

**Supplementary Table 9. Comparison of CpG methylation frequencies from whole genome bisulfite sequencing (WGBS) Illumina data with Cas9-targeted Nanopore data for each method using single score thresholds obtained by the maximum value of (TPR-FPR).** We used the score cutoffs that maximized TPR-FPR in the mixture dataset 1 (Supplementary Table 3). For each method we provide the number of sites (N), the Pearson's correlation (r), coefficient of determination ($r^2$), the Spearman's rank correlation (ρ), and the root mean square error (RMSE) for the comparison of the percentage methylation predicted from Nanopore with the percentage methylation calculated from whole genome bisulfite sequencing (WGBS) data. We show the results for five tested tools and METEORE combining DeepSignal and Megalodon using a random forest (RF) (parameters: max_depth=3 and n_estimator=10) or a regression (REG) model.

| | *N* | *r* | *$r^2$* | *ρ* | *RMSE* |
|---|---|---|---|---|---|
| Nanopolish | 2171 | 0.9265 | 0.8584 | 0.7943 | 0.2061 |
| DeepSignal | 2171 | 0.9654 | 0.9319 | 0.8149 | 0.1246 |
| Megalodon | 2171 | 0.9602 | 0.9219 | 0.8195 | 0.1372 |
| Tombo | 2171 | 0.8936 | 0.7986 | 0.7802 | 0.2343 |
| Guppy | 2171 | 0.9034 | 0.8162 | 0.7957 | 0.2148 |
| METEORE (RF) | 2171 | 0.9668 | 0.9348 | 0.8262 | 0.1221 |
| METEORE (REG) | 2171 | 0.9646 | 0.9305 | 0.8263 | 0.1271 |

**Supplementary Table 10. Comparison of CpG methylation frequencies from whole genome bisulfite sequencing (WGBS) Illumina data with Cas9-targeted Nanopore data from Gilpatrick et al. 2020 for each method using single score thresholds obtained by the maximum value of (TPR-FPR).** We used the score cutoffs that maximized TPR-FPR in the mixture dataset 1 (Supplementary Table 3). For each method we provide the number of sites (N), the Pearson's correlation (r), coefficient of determination ($r^2$), the Spearman's rank correlation (ρ), and the root mean square error (RMSE) for the comparison of the percentage methylation predicted from Nanopore with the percentage methylation calculated from whole genome bisulfite sequencing (WGBS) data. We show the results for five tested tools and METEORE combining DeepSignal and Megalodon using a random forest (RF) (parameters: max_depth=3 and n_estimator=10) or a regression (REG) model.

| | *N* | *r* | $r^2$ | *ρ* | *RMSE* |
|---|---|---|---|---|---|
| Nanopolish | 1621 | 0.8726 | 0.7614 | 0.8327 | 0.2165 |
| DeepSignal | 1731 | 0.9220 | 0.8500 | 0.8805 | 0.1698 |
| Megalodon | 1723 | 0.8786 | 0.7720 | 0.8643 | 0.2336 |
| Tombo | 1733 | 0.8150 | 0.6642 | 0.7924 | 0.2468 |
| Guppy | 1733 | 0.7345 | 0.5394 | 0.7275 | 0.3243 |
| METEORE (RF) | 1723 | 0.9185 | 0.8437 | 0.8874 | 0.1812 |
| METEORE (REG) | 1722 | 0.9167 | 0.8404 | 0.8917 | 0.1866 |

**Supplementary Table 11. Comparison of CpG methylation frequencies from whole genome bisulfite sequencing (WGBS) Illumina data with Cas9-targeted Nanopore data for each method using the double cutoff obtained by discarding 10% of reads.** For each site, we removed the 10% of reads with scores closest to the cross point of the FPR and 1-TPR curves in the mixture dataset 1 (Supplementary Table 4). For each method we provide the number of sites (N), the Pearson's correlation (r), coefficient of determination ($r^2$), the Spearman's rank correlation (ρ), and the root mean square error (RMSE) for the comparison of the percentage methylation predicted from Nanopore with the percentage methylation calculated from whole genome bisulfite sequencing (WGBS) data. METEORE is the combination model with the adjusted parameters of a random forest (max_depth=3 and n_estimator=10) combining DeepSignal and Megalodon.

| | *N* | *r* | $r^2$ | *ρ* | *RMSE* |
|---|---|---|---|---|---|
| Nanopolish | 2168 | 0.9376 | 0.8792 | 0.8128 | 0.1641 |
| DeepSignal | 2171 | 0.9677 | 0.9364 | 0.8176 | 0.1187 |
| Megalodon | 2171 | 0.9492 | 0.901 | 0.8128 | 0.1632 |
| Tombo | 2171 | 0.9013 | 0.8123 | 0.7809 | 0.2177 |
| Guppy | 2171 | 0.8867 | 0.7862 | 0.7811 | 0.2248 |
| METEORE (RF) | 2171 | 0.9652 | 0.9317 | 0.8274 | 0.1276 |
| METEORE (REG) | 2171 | 0.9653 | 0.9318 | 0.8310 | 0.1253 |

**Supplementary Table 12. Comparison of CpG methylation frequencies from whole genome bisulfite sequencing (WGBS) Illumina data with Cas9-targeted Nanopore data from Gilpatrick et al. 2020 for each method using the double cutoff obtained by discarding 10% of reads.** For each site, we removed the 10% of reads with scores closest to the cross point of the FPR and 1-TPR curves in the mixture dataset 1 (Supplementary Table 4). For each method we provide the number of sites (N), the Pearson's correlation (r), coefficient of determination ($r^2$), the Spearman's rank correlation (ρ), and the root mean square error (RMSE) for the comparison of the percentage methylation predicted from Nanopore with the percentage methylation calculated from WGBS data. We show the results for five tested tools and METEORE combining DeepSignal and Megalodon using a random forest (RF) (parameters: max_depth=3 and n_estimator=10) or a regression (REG) model.

| | No. of CPUs used | Real time per CPU (min) | Peak memory (GB) | Bases per second |
|---|---|---|---|---|
| Nanopolish | 2 | 10.2 | 0.2 | 25433 |
| DeepSignal | 9 | 334.8 | 23.7 | 775 |
| Tombo | 9 | 30.4 | 23.6 | 8533 |
| Megalodon | 11 | 3258.7 | 1.4 | 80 |
| Megalodon (GPU) | 1 GPU | 3.8 per GPU | 2.4 | 69710 |
| Guppy | 11 | 1494.6 | 3.3 | 174 |
| Guppy (GPU) | 1 GPU | 7.0 per GPU | 0.6 | 37596 |
| DeepMod | 8 | 176.0 | 2.5 | 1474 |
| METEORE (RF) | 1 | 0.1 | 0.3 | - |
| METEORE (REG) | 1 | 1.8 | 9.5 | - |

**Supplementary Table 13. Runtime and memory usage for each tested tool.** We tested each pipeline on the m50 set (2,432 reads and a total of 15,564,827 bases) from the mixture dataset 1. We recorded the real time (wall clock time) from start to finish of the pipeline/command(s) for all tested tools except for METEORE, which took a fast5 directory as an input and output a prediction at genome level (i.e., methylation frequency for each site). For METEORE, we only recorded the time needed for running a Python script to make the per-read consensus predictions. We used METEORE combining DeepSignal and Megalodon using a random forest (RF) (parameters: max_depth=3 and n_estimator=10) and a regression (REG) model, where both models took per-read prediction outputs generated from the selected tools. We ran each tested tool on a computer with 12 CPU processors (Intel Core i7 (8th Gen) 8700 @ 3.2 GHz). Guppy can be run on CPUs or GPUs, but on a GPU the basecalling speed increase significantly. Megalodon requires Guppy for the basecalling step and uses GPU-enabled Guppy by default. If Megalodon is used with Guppy (CPU), the --guppy-timeout argument should be specified (here we used 200 seconds) to allow sufficient time for calling a read during CPU basecalling. Additionally, we ran Megalodon and Guppy on another computer with a GPU (GeForce RTX 2080 Ti) and 32 CPU processors (AMD Ryzen threadripper 2950x). Real time per CPU = real (wall-clock) time taken x no. of CPUs. Bases per second = total no. of bases/(real time per CPU x 60).
