## Supplementary material for "Systematic benchmarking of tools for CpG methylation detection from Nanopore sequencing": Description of the supplementary data files

### Description of Additional Supplementary Files

**Supplementary Data 1**

Filename: Supplementary_data_1.tsv

Description: 100 CpG sites used in mixture dataset 1. A tab separated file including the following fields:

1. Chromosome
2. Ending position

**Supplementary Data 2**

Filename: Supplementary_data_2.tsv

Description: 50 CpG sites used in mixture dataset 2. A tab separated file including the following fields:

1. Chromosome
2. Ending position

**Supplementary Data 3**

Filename: Supplementary_data_3.tsv

Description: Fast5 files used to generate each subset in mixture dataset 1. The fast5 files were generated by Simpson et al. (2017) and can be downloaded from the European Nucleotide Archive (ENA) under accession PRJEB13021 (ERR1676719 for negative control and ERR1676720 for positive control). A tab separated file including the following fields:

1. Name of the subset (m0, m10, m20…)
2. Fast5 filename

**Supplementary Data 4**

Filename: Supplementary_data_4.tsv

Description: Fast5 files used to generate each subset in mixture dataset 2. The fast5 files was generated by Simpson et al. (2017) and can be downloaded from the European Nucleotide Archive (ENA) under accession PRJEB13021 (ERR1676719 for negative control and

ERR1676720 for positive control). The reads used in mixture dataset 2 are different from mixture dataset 1. A tab separated file including the following fields:

1. Name of the subset (m0, m10, m20…)
2. Fast5 filename

**Supplementary Data 5**

Filename: Supplementary_data_5.xlsx

Description: Methylation frequencies of CpG sites calculated by each tested tool for each subset in control mixture dataset 1. An Excel file including the following fields:

1. Chromosome
2. Ending position
3. % Methylated sets
4. Expected methylation frequency
5. Predicted methylation frequency
6. Method
7. Left bound of each 10% methylated bin
8. Right bound of each 10% methylated bin
9. Predicted [inside/outside] a 10% window around the expected methylation proportion

**Supplementary Data 6**

Filename: Supplementary_data_6.xlsx

Description: The number of true negatives for each tested tool according to different thresholds for the predicted methylation frequency below which a site was called unmethylated, using 0% methylated set from mixture dataset 1. An Excel file including the following fields:

1. Threshold
2. Method
3. Number of true negatives

**Supplementary Data 7**

Filename: Supplementary_data_7.xlsx

Description: The number of true negatives for each tested tool according to different thresholds for the predicted methylation frequency above which a site was called methylated, using 100% methylated set from mixture dataset 1. An Excel file including the following fields:

1. Threshold
2. Method
3. Number of true negatives

**Supplementary Data 8**

Filename: Supplementary_data_8.xlsx

Description: Per-read performance of a binary classifier at all possible classification thresholds for five tested tools measured by receiver operating characteristic (ROC) curve, using reads from 0% methylated set from mixture dataset 1. An Excel file including the following fields:

1. Threshold
2. Specificity, i.e. TN/(TN +FP) = 1−FPR
3. Sensitivity, i.e. TP/(TP/FN) = 1−FNR = TPR
4. FPR
5. Method

, where TP = true positives, FP = false positives, TN = true negatives, FN = false negatives, TPR = true positive rate, and FPR = false positive rate.

**Supplementary Data 9**

Filename: Supplementary_data_9.xlsx

Description: Per-read performance of a binary classifier at all possible classification thresholds for five tested tools measure by precision-recall (PR) curve, using reads from 0% methylated set from mixture dataset 1. An Excel file including the following fields:

1. Threshold
2. Recall, i.e. TP/(TP/FN) = 1−FNR = TPR

3. Precision, i.e. TP/(TP+FP) = 1−FDR
4. Method

, where TP = true positives, FP = false positives, TN = true negatives, FN = false negatives, TPR = true positive rate, and FDR = false discovery rate.

**Supplementary Data 10**

Filename: Supplementary_data_10.xlsx

Description: Methylation frequencies of CpG sites calculated by each tested tool for each subset in control mixture dataset 2. An Excel file including the following fields:

1. Chromosome
2. Ending position
3. % Methylated sets
4. Expected methylation frequency
5. Predicted methylation frequency
6. Method
7. Left bound of each 10% methylated bin
8. Right bound of each 10% methylated bin
9. Predicted [inside/outside] a 10% window around the expected methylation proportion

**Supplementary Data 11**

Filename: Supplementary_data_11.xlsx

Description: Methylation frequencies and coverage of CpG sites reported by each tested tool across the 10 targeted regions from our Cas9-targeted Nanopore sequencing (nCATS) data. An Excel file including the following fields:

1. Chromosome
2. Ending position
3. Methylation frequency reported by Nanopolish
4. Coverage reported by Nanopolish
5. Methylation frequency reported by DeepSignal
6. Coverage reported by DeepSignal

7. Methylation frequency reported by Megalodon
8. Coverage reported by Megalodon
9. Methylation frequency reported by Tombo
10. Coverage reported by Tombo
11. Methylation frequency reported by Guppy
12. Coverage reported by Guppy
13. Methylation frequency reported by DeepMod
14. Coverage reported by DeepMod
15. Methylation frequency reported by METEORE (RF)
16. Coverage reported by METEORE (RF)
17. Methylation frequency reported by METEORE (REG)
18. Coverage reported by METEORE (REG)
19. Methylation frequency reported by whole genome bisulfite sequencing (WGBS)
20. Depth of coverage reported from the BAM file
21. WGBS methylation bin
22. Target region

**Supplementary Data 12**

Filename: Supplementary_data_12.xlsx

Description: Methylation frequencies and coverage of CpG sites reported by each tested tool across the 8 targeted regions from the Cas9-targeted Nanopore sequencing (nCATS) data generated by Gilpatrick et al. (2020). An Excel file including the following fields:

1. Chromosome
2. Ending position
3. Methylation frequency reported by Nanopolish
4. Coverage reported by Nanopolish
5. Methylation frequency reported by DeepSignal
6. Coverage reported by DeepSignal
7. Methylation frequency reported by Megalodon
8. Coverage reported by Megalodon
9. Methylation frequency reported by Tombo

10. Coverage reported by Tombo
11. Methylation frequency reported by Guppy
12. Coverage reported by Guppy
13. Methylation frequency reported by DeepMod
14. Coverage reported by DeepMod
15. Methylation frequency reported by METEORE (RF)
16. Coverage reported by METEORE (RF)
17. Methylation frequency reported by METEORE (REG)
18. Coverage reported by METEORE (REG)
19. Methylation frequency reported by whole genome bisulfite sequencing (WGBS)
20. Depth of coverage reported from the BAM file
21. WGBS methylation bin
22. Target region
